# Multiomic interpretation of fungus-infected ant metabolomes during manipulated summit disease

**DOI:** 10.1101/2023.01.19.524761

**Authors:** I. Will, G. M. Attardo, C. de Bekker

## Abstract

*Camponotus floridanus* ants show altered behaviors followed by a fatal summiting phenotype when infected with manipulating *Ophiocordyceps camponoti-floridani* fungi. Host summiting as a strategy to increase transmission is also observed with parasite taxa beyond fungi, including aquatic and terrestrial helminths and baculoviruses. The drastic phenotypic changes can sometimes reflect significant physiological changes within host cells that span molecular levels from metabolites to nucleic acids. Nevertheless, the underlying mechanisms still need to be fully characterized. To investigate the small molecules producing summiting behavior, we infected *C. floridanus* ants with *O. camponoti-floridani* and sampled their heads for LC-MS/MS when we observed the characteristic summiting phenotype. We link this metabolomic data with our previous genomic and transcriptomic data to propose mechanisms that underlie manipulated summiting behavior in “zombie ants.” This “multiomic” evidence points toward the dysregulation of neurotransmitter levels and neuronal signaling. We propose that these processes are altered during infection and manipulation based on 1) differential expression of neurotransmitter synthesis and receptor genes, 2) altered abundance of metabolites and neurotransmitters (or their precursors) with known behavioral effects in ants and other insects, and 3) possible suppression of a connected immunity pathway. We additionally report signals for metabolic activity during manipulation related to primary metabolism, detoxification, and anti-stress protectants.

## Introduction

Infections often lead to altered host phenotypes that can be adaptive for the host or parasite. Parasite-adaptive changes in animal host behavior are mostly referred to as manipulations. Apparent manipulations of host behavior and physiology occur in a broad range of host and parasite taxa. While descriptive examples are plenty, the underlying mechanisms are still in the process of being revealed ^1–5^.

The ant-manipulating *Ophiocordyceps* (zombie ant fungi) have been leveraged in laboratory studies to better understand the mechanistic basis of behavioral manipulation ^5–7^. Infected ants display summit disease: a common parasitic manipulation observed in invertebrates where the manipulated individual occupies an elevated location and dies stereotypically ^5, 8^. These final, elevated, exposed summit positions can be adaptive for the parasite by either aiding trophic transmission or direct dispersal of infectious propagules. In the case of *Ophiocordyceps*, ants latch and bite onto their final summit perches until death, which is thought to promote development of the fungal fruiting body and dispersal of the spores therein ^9–13^. Moreover, preceding this fatal change in behavior, infected ants also deviate from foraging trails and increase locomotor activity, reduce nestmate communication, and convulse ^12–17^.

These various changes may represent behaviors that can be easily coopted for manipulation by the fungal parasite ^5^. As such, the parasite may be taking advantage of existing host processes and symptoms without relying on costly mechanisms to “rewire” their hosts ^8, 18^. A profound understanding of the fitness costs and benefits of these behaviors, for both the host and parasite, may reveal that some could also be adaptive for the host, at the individual or group level (perhaps especially so for eusocial animals, such as ants). Regardless of the ultimate “source” of the altered behaviors observed, physiological and molecular characterizations of hosts displaying modified behavior may offer a glimpse into how parasites (or hosts) modulate animal behavior, the coevolution of specialized host-parasite relationships, and undescribed properties of proteins and metabolites involved.

In the myrmecophilous *Ophiocordyceps*, investigations of phenotypes, genomes, transcriptomes, and metabolomes continue to bring various mechanistic hypotheses of behavioral manipulation into focus. The range of hypotheses comprise fungal neuromodulators and effector proteins, host circadian rhythms and phototropism, insect hormones, fungal neuroprotectants, and ant muscular hyperactivity ^6, 7, 14, 19–24^. Previous metabolomic studies, centered on the *Ophiocordyceps kimflemingiae* zombie ant fungus and its host *Camponotus castaneus*, have identified various metabolites associated with manipulation. These metabolomic signals also varied in a species-specific manner, changing when either the parasite or host was exchanged for a different species ^21, 22, 25^. However, these studies did not place the results within the context of other types of “omics” data. Doing so would present the opportunity to distinguish the most robust lines of evidence from the more stochastic signals that might differ across experimental conditions and setups.

Here, we present a “multiomic” approach by conducting a metabolomic analysis of *Camponotus floridanus* (Florida carpenter ant) manipulated by *Ophiocordyceps camponoti-floridani* (Florida zombie ant fungus) interpreted in the context of our previous transcriptomic and genomic data ^24^. To add to our existing transcriptomic study, we collected metabolomic data using three liquid chromatography-tandem mass spectrometry (LC-MS/MS) protocols. Our study compares whole head samples of healthy *C. floridanus* ants and ants manipulated by *O. camponoti-floridani*, allowing direct comparison to the transcriptomic ant-head data. Whole heads contain the chemical profiles of ant neural tissue, muscle, and hemolymph in addition to fungal cells that all may contain metabolites of most interest ^13, 15, 20–22, 25, 26^.

We primarily focused our interpretation and discussion of the results on altered metabolic pathways that are supported by a combination of metabolite and gene signals. This improves mechanistic interrogation as the key processes of manipulation could involve specialized and general mechanisms, host and parasite effectors, and changes from the genetic to chemical level. At the same time, combining multiple omics approaches and datasets allowed us to identify consistent and robust signals for key biological processes. To analyze our metabolomics data we also combined multiple statistical methods to reduce the inherent bias that any given analysis introduces. Data that represent substantial biological changes are more likely to be selected by multiple complementary tests. Our reporting, therefore, highlights manipulation-associated metabolite features that were selected by multiple statistical methods. With this, we propose multiple pathways and compounds that reflect, or drive, manipulated behavior in *Ophiocordyceps* infected ants.

## Results & discussion

### Infection mortality, observations of manipulation, and LC-MS/MS

Similar to previous laboratory infections with *O. camponoti-floridani* ^24^, we collected *C. floridanus* displaying manipulated summiting between four hours before (zeitgeber time, ZT 20) to half an hour after dawn (ZT 0.5), beginning three weeks after infection (Fig. 1). Sham-treated healthy ants showed no mortality until 29 days post injection (dpi), whereas the survival rate among infected ants gradually decreased from the beginning of the infection experiment, with a steeper drop during the first half of the manipulation phase starting in week three. As such, survival was markedly different between treatments (p = 4.1E-7, log-rank test) (Fig. 1A). In total, three sham-treated ants died out of the 28 (11%) that survived the initial injection procedure. Twenty-three infected ants were manipulated out of the 39 (59%) that survived the injection. The remaining 16 infected ants died with fungal infection, indicated by fungal blastospores present upon microscopic investigation, but without observed behavioral manipulation. From these experimental infections, we sampled 20 manipulated ants and 20 healthy (sham) ants for LC-MS/MS. We also produced three blank samples to contrast our treatment samples against. Using these samples, we produced three LC-MS/MS datasets: “biogenic amines” (2,955 features), “polyphenols” (4,315 features), and “lipids” (4,924 features).

**Figure 1.**
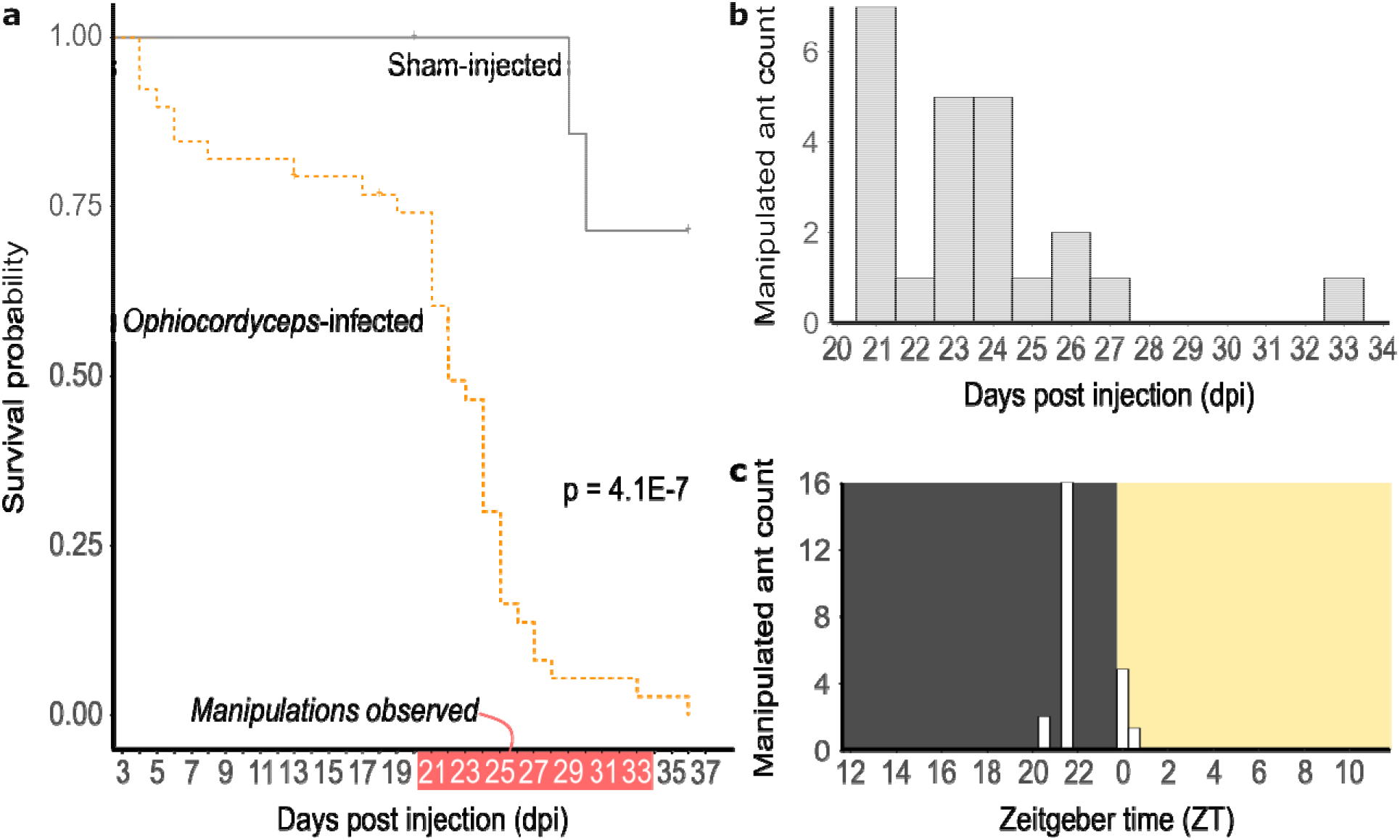
Ant manipulations occur at stereotypical days and times. **a**) *Ophiocordyceps camponoti-floridani* infection leads to significantly increased mortality compared to sham treatment (p = 4.1E-7, log-rank test). **b**) Manipulated ants most often appeared 21-27 dpi, but as late as 33 dpi. **c**) Summiting behavior always occurred around dawn, most often 2.5 h before lights turned on, at ZT 21.5.

### Metabolomic feature selection and grouping for pathway enrichment analyses

We combined multiple analyses to stringently select “top features” and metabolic pathways for further biological interpretation. We identified metabolite features that distinguished healthy and manipulated ants by their *i*) being differentially abundant metabolites (DAMs), *ii*) loading values in principal component analyses (PCAs), or *iii*) importance for navigating decision trees (Boruta) ^27^ (Fig. 2A). Combining all three LC-MS/MS datasets, approximately 60% of features were DAMs, with most DAMs increasing in abundance in manipulated ants compared to healthy controls (Fig. 2B, Supplementary Table S1). Separation of treatment groups by PCA only clearly clustered along PC1 (ca. 60% to 80%), with PC2 (ca. 4% to 9%) and other PCs (not shown) describing little of the variation between treatments (Fig. 2C). Using Boruta random forests, we found ∼40% of features to be “confirmed” as “important” for distinguishing treatment types (Supplementary Table S1). We performed pathway analyses with top features that satisfied at least two of the following criteria: the feature was *i*) found to be a DAM, *ii*) among the top 50% of absolute PC1 loading values, or *iii*) confirmed by Boruta. Most top features that satisfied at least two requirements (n = 6,468 features, 53% of a total 12,194 features), in fact satisfied all three (n = 4,656, 72% of features passing two criteria) (Fig. 2D, Supplementary Table S1, Supplementary Data S1). The feature set passing at least two filters contained 347 of the 760 (46%) features that were functionally annotated. This many features passing our stringent selection approach is likely due to a combination of small molecules related specifically to infection processes and general aspects of fungal metabolism, as *O. camponoti-floridani* cells were only present in manipulated ants.

**Figure 2.**
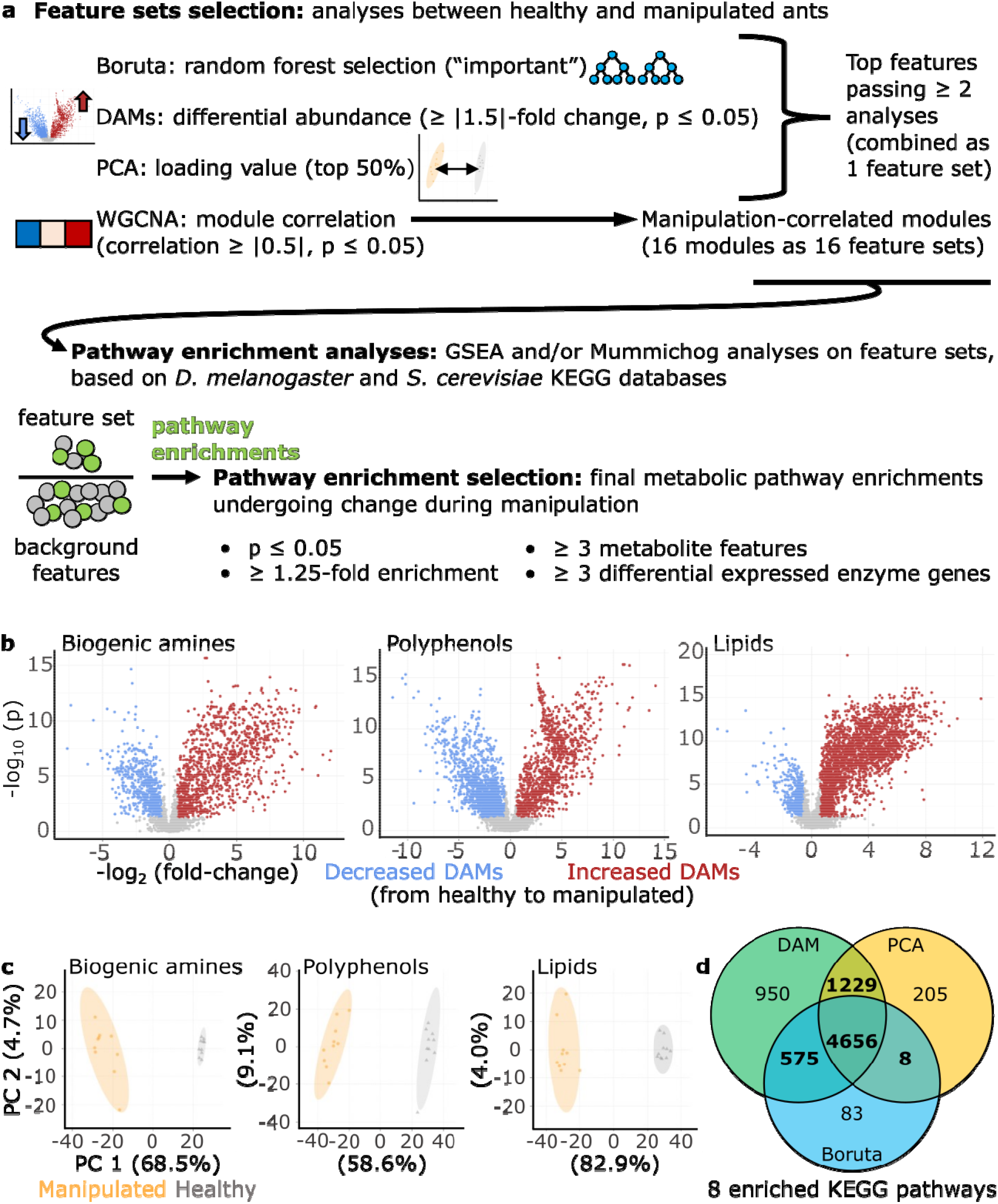
Selection of top features based on three independent analyses. **a**) We used three independent statistical tests to select a single top features for metabolic pathway enrichment analyses. Additionally, we used a weighted co-abundance network approach (analogous to WGCNA used for gene expression data) to form manipulation-correlated feature modules (resulting in 16 feature sets) for enrichment analyses. Results from those enrichment analyses were subsequently subjected to multiple filters to arrive at a final set of pathways and feature sets of interest. We also performed an enrichment analysis on a set of features only present in manipulated samples, but we found no overrepresented pathways and this feature set is not depicted in the figure. **b**) Volcano plots per dataset show DAMs that increased (red) or decreased (blue) at manipulation according to a 1.5-fold change and p ≤ 0.05 t-test threshold. **c**) PCAs for each dataset show distinct clustering of samples between manipulated (orange) and healthy ants (gray) along PC1. d) Features that passed at least two selection criteria (bold) were classified as top features and used for further pathway analyses. Top features were enriched for eight KEGG metabolic pathways.

We analyzed the 6,468 top features with Mummichog and a gene set enrichment analysis (GSEA)-based approach to detect enriched Kyoto Encyclopedia of Genes and Genomes (KEGG) metabolic pathways (Fig. 2A) ^28–30^. We found eight significantly enriched pathways that passed our p- value, fold-enrichment, metabolite feature count, and enzyme differentially expressed gene (DEG) count thresholds (Fig. 2A, Fig. 2D, Supplementary Data S2). These enzyme DEGs were identified by their association with KEGG metabolic pathways and the previously published expression data from our *Ophiocordyceps*-infection study in which the heads of manipulated ants were compared to those of healthy ants using RNASeq ^24^. Combining enriched pathways from these top features and the network module analysis (see below) (Fig. 2A, Fig. 3, Table 1), most enzyme DEGs were downregulated both in the ant (90%) and the fungus (62%). This degree of downregulation is higher than previously found for transcriptome-wide DEGs during manipulation ^24^ (Supplementary Discussion S1). We infer a positive correlation between DEG transcription and protein levels as the most parsimonious interpretation of the data. Although, gene expression reflecting homeostatic responses to changes in functional protein abundance or a lack of consistent correlation are alternative explanations, which future proteomic data could help make clear.

**Figure 3.**
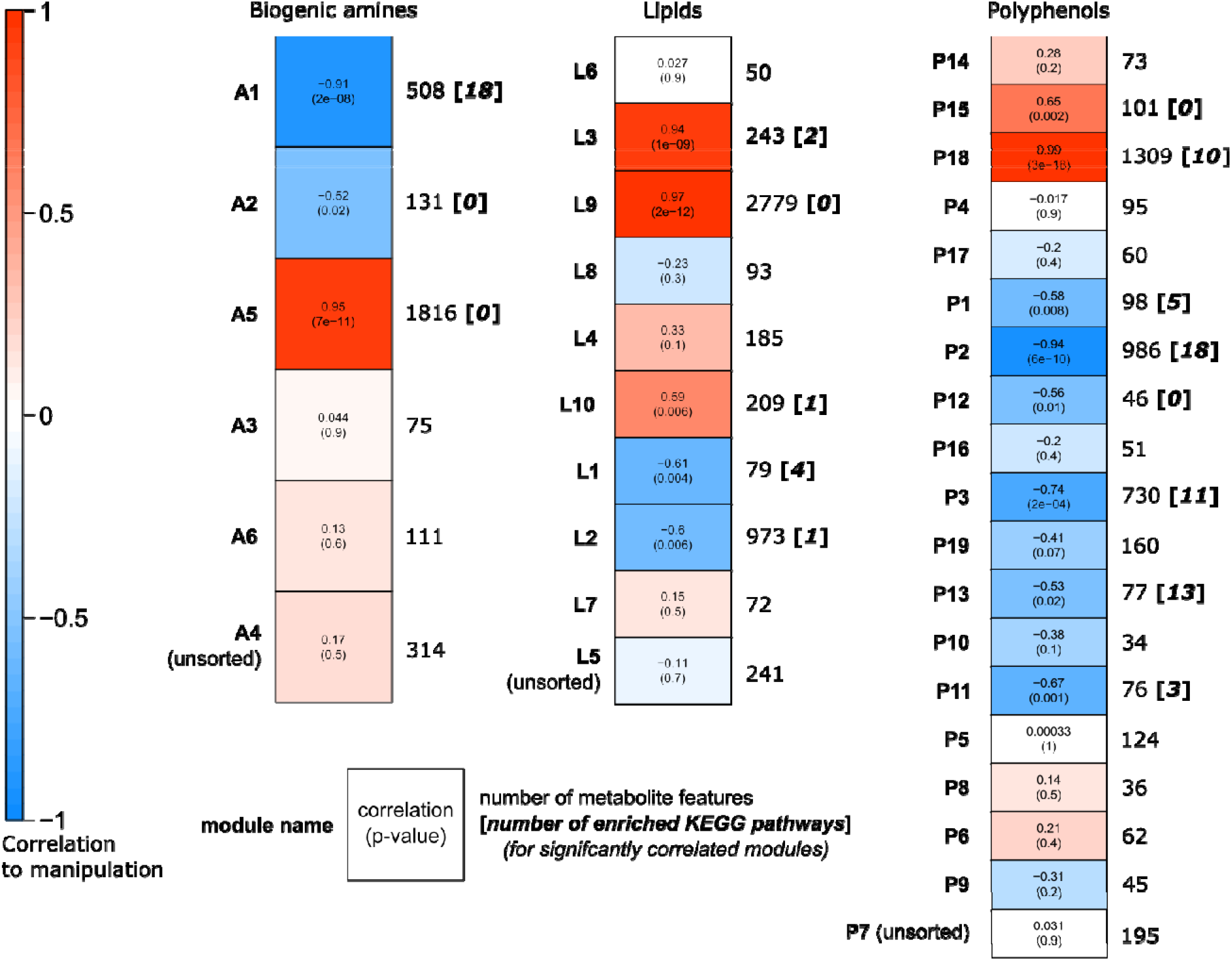
WGCNA network modules of metabolite features, their correlation to manipulation, and enriched pathways. In total we detected three, five, and eight network modules that were significantly correlated to manipulation (Student correlation ≥ |0.5|, p ≤ 0.05) in the biogenic amines, lipids, and polyphenol datasets, respectively. “Unsorted” modules contain all features that were not successfully correlated to other features in a network.

**Table 1.**
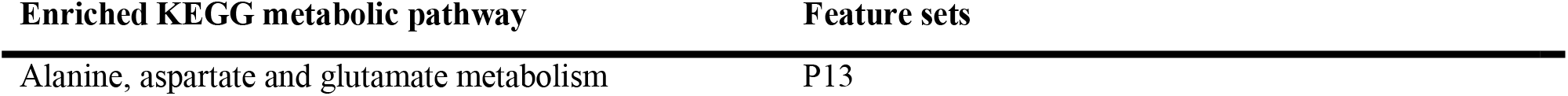
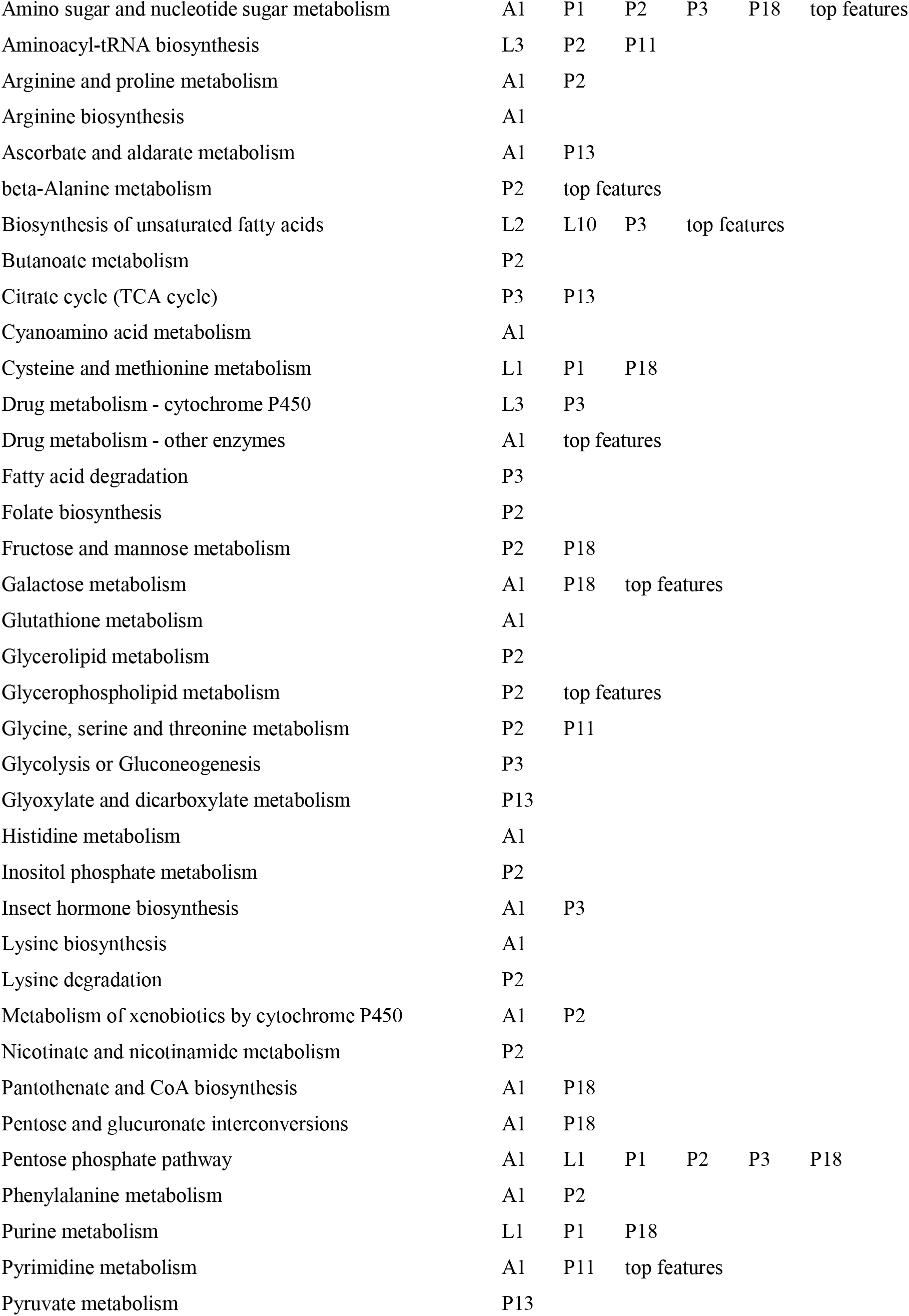
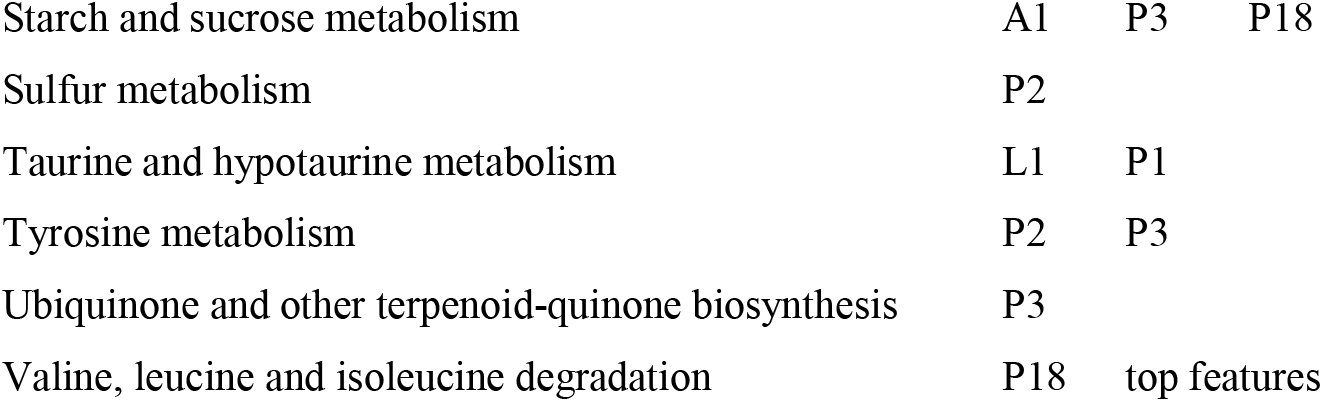
Enriched KEGG pathways selected across analyzed feature sets. For KEGG pathways that were found in Mummichog or GSEA analyses (p ≤ 0.05) to be of interest, we also required them to show a minimum 1.25-fold enrichment, have at least three features contributing to that enrichment, and the previous transcriptomics dataset ^24^ to contain at least three DEGs homologous to representative enzymes within that pathway. Pathways are listed alphabetically with all feature sets (network modules or the top features set) that were enriched for that pathway. Network modules names: biogenic amine (A1-A6), lipid (L1-L10), and polyphenol (P1-P19). Only 16 of these 35 modules were correlated to manipulation and appear in this table.

| Enriched KEGG metabolic pathway | Feature sets |
| --- | --- |
| Alanine, aspartate and glutamate metabolism | P13 |

|  |  |  |  |  |  |  |
| --- | --- | --- | --- | --- | --- | --- |
| Amino sugar and nucleotide sugar metabolism | A1 | P1 | P2 | P3 | P18 | top features |
| Aminoacyl-tRNA biosynthesis | L3 | P2 | P11 |  |  |  |
| Arginine and proline metabolism | A1 | P2 |  |  |  |  |
| Arginine biosynthesis | A1 |  |  |  |  |  |
| Ascorbate and aldarate metabolism | A1 | P13 |  |  |  |  |
| beta-Alanine metabolism | P2 | top features |  |  |  |  |
| Biosynthesis of unsaturated fatty acids | L2 | L10 | P3 | top features |  |  |
| Butanoate metabolism | P2 |  |  |  |  |  |
| Citrate cycle (TCA cycle) | P3 | P13 |  |  |  |  |
| Cyanoamino acid metabolism | A1 |  |  |  |  |  |
| Cysteine and methionine metabolism | L1 | P1 | P18 |  |  |  |
| Drug metabolism - cytochrome P450 | L3 | P3 |  |  |  |  |
| Drug metabolism - other enzymes | A1 | top features |  |  |  |  |
| Fatty acid degradation | P3 |  |  |  |  |  |
| Folate biosynthesis | P2 |  |  |  |  |  |
| Fructose and mannose metabolism | P2 | P18 |  |  |  |  |
| Galactose metabolism | A1 | P18 | top features |  |  |  |
| Glutathione metabolism | A1 |  |  |  |  |  |
| Glycerolipid metabolism | P2 |  |  |  |  |  |
| Glycerophospholipid metabolism | P2 | top features |  |  |  |  |
| Glycine, serine and threonine metabolism | P2 | P11 |  |  |  |  |
| Glycolysis or Gluconeogenesis | P3 |  |  |  |  |  |
| Glyoxylate and dicarboxylate metabolism | P13 |  |  |  |  |  |
| Histidine metabolism | A1 |  |  |  |  |  |
| Inositol phosphate metabolism | P2 |  |  |  |  |  |
| Insect hormone biosynthesis | A1 | P3 |  |  |  |  |
| Lysine biosynthesis | A1 |  |  |  |  |  |
| Lysine degradation | P2 |  |  |  |  |  |
| Metabolism of xenobiotics by cytochrome P450 | A1 | P2 |  |  |  |  |
| Nicotinate and nicotinamide metabolism | P2 |  |  |  |  |  |
| Pantothenate and CoA biosynthesis | A1 | P18 |  |  |  |  |
| Pentose and glucuronate interconversions | A1 | P18 |  |  |  |  |
| Pentose phosphate pathway | A1 | L1 | P1 | P2 | P3 | P18 |
| Phenylalanine metabolism | A1 | P2 |  |  |  |  |
| Purine metabolism | L1 | P1 | P18 |  |  |  |
| Pyrimidine metabolism | A1 | P11 | top features |  |  |  |
| Pyruvate metabolism | P13 |  |  |  |  |  |

|  |  |  |  |
| --- | --- | --- | --- |
| Starch and sucrose metabolism | A1 | P3 | P18 |
| Sulfur metabolism | P2 |  |  |
| Taurine and hypotaurine metabolism | L1 | P1 |  |
| Tyrosine metabolism | P2 | P3 |  |
| Ubiquinone and other terpenoid-quinone biosynthesis | P3 |  |  |
| Valine, leucine and isoleucine degradation | P18 | top features |  |

The eight KEGG metabolic pathways enriched among top features included signals that suggest altered primary metabolism related to amino sugars, galactose, valine and leucine, pyrimidines, and fatty acids (Table 1). Changes in primary metabolism could reflect various processes, including diseased host metabolism, parasite presence or acquisition of host resources, or host-parasite cross-talk ^31–37^. Furthermore, the previous transcriptomics data prompted hypotheses connecting behavioral manipulation to nutritional state and insulin-related signaling ^24^.

Other enriched pathways plausibly reflect manipulated behavior and infection in different aspects of metabolism (Table 1). We found KEGG pathway “drug metabolism – other enzymes” enriched among top features. Broadly suggesting antagonistic host-parasite chemical interactions, this pathway annotation contains multiple sub-pathways corresponding to different toxic metabolites, which are difficult to resolve as we lacked positive identification of these specific compounds in our data (Supplementary Discussion S2). Additionally, as aspects of enriched “beta-alanine metabolism,” we detected an apparent carnosine feature and its constituent amino acids as DAMs (Supplementary Discussion S3). Although we lack robust transcriptomic support for the synthesis of carnosine (Supplementary Discussion S3), the combination of pathway enrichment and metabolite feature data indicated that carnosine is produced from host amino acids during infection. This compound may act as a cytoprotectant during the stresses behavioral manipulation ^38–41^. We also further discuss enriched “glycerophospholipid metabolism” in context with its multiomic signals and identified metabolites in detail below.

Many metabolite features in our dataset were uniquely represented in manipulated samples: 454 features were never detected in healthy sham treatment ants and 1,781 additional features either were detected in less than 50% of the healthy ant samples and/or their median peak value was less than two-fold that of blank controls (2,235 features total). These features may represent uniquely produced fungal molecules or ant molecules that are tightly upregulated with *Ophiocordyceps*-infection while being scarce in healthy ants. Although we predicted that hallmarks of infection and manipulation processes would be overrepresented among these unique features, we did not find any significant pathway enrichments. This could indicate that these metabolites are not linked by common metabolic pathways and/or many of these metabolites were not well annotated in the KEGG databases (e.g., species-specific metabolites).

Co-expression (or co-abundance) networks can suggest linked biological functions and activity in the same metabolic processes ^42–45^. To cluster metabolite features that similarly change abundance between healthy and manipulated *C. floridanus* treatments, we used a weighted gene co-expression network analysis (WGCNA) based approach (Fig. 2A, Fig. 3) ^46^. This resulted in a total of six biogenic amine (A1-A6), 10 lipid (L1-L10), and 19 polyphenol (P1-P19) modules. Of these network modules, three biogenic amine, five lipid, and eight polyphenol modules were significantly correlated to *Ophiocordyceps* manipulation (Fig. 3, Table 1). Due to our sparse annotation data obtained from mass spectrometry database matching, we were not able to consistently compare hub features. Instead, we considered all available annotations within a network module and Mummichog pathway enrichments (Fig. 2A). Across all network modules significantly correlated to the transition from healthy to manipulated ants, we found 44 unique enriched KEGG pathways that passed our selection criteria, which included the eight found from the top features set (Table 1, Supplementary Data S2).

### Biogenic monoamine neurotransmitters as putative proximate drivers of *Ophiocordyceps* infection-related behaviors

A combination of pathway enrichments, annotated DAMs, and DEGs suggest that shifts in metabolism associated with neurotransmitters relate to behavioral alterations in *C. floridanus* infected and manipulated by *O. camponoti-floridani*. The monoamine neurotransmitters dopamine, serotonin, and octopamine (analogous to vertebrate norepinephrine) have been implicated in modulating locomotor, foraging, learning, social, reproductive, and aggressive behaviors in many insects, and have been tested in social insects, including ants ^47–51^. Serotonin, octopamine, and melanin derived from L-3,4-dihydroxyphenylalanine (L-DOPA), or tyrosine and dopamine, have roles in insect immunity as well ^52, 53^. Although we did not directly identify dopamine, serotonin, or octopamine in our metabolomic data, we did measure features annotated as biosynthetically related compounds. Furthermore, gene expression networks negatively correlated between the ant and fungus were enriched for host neuronal function and parasite manipulation-associated secreted proteins. This correlation suggested that parasite effectors promote, directly or indirectly, the downregulation of host gene modules related to neuronal maintenance, circadian rhythm, olfaction, and memory ^24^.

Moreover, these neurotransmitters can affect behaviors relatable to *Ophiocordyceps* disease phenotypes. In *Formica polyctena* wood ants and *Odontomachus kuroiwae* trap-jaw ants, administration of dopamine and serotonin induced aggressive mandible-opening or biting behaviors and defensive responses in the laboratory ^54, 55^. In the field, *Pogonomyrmex barbatus* harvester ant dopamine levels positively correlated with foraging trips ^56^. Similarly, pharmacological depletion of serotonin led to reduced and impaired trail-following foraging behavior in *Pheidole dentata* ants ^57^. In *Formica aquilonia*, there is evidence for octopamine increasing aggressive behaviors ^58^. However, increased aggression upon octopamine treatment was not found in *F. polyctena* ^54^. Octopamine additionally appears to modulate the social behavior of trophallaxis in *Camponotus fellah*, while serotonin had no significant effect ^59^. As such, the exact effects of these neurotransmitters can differ by species and social or physiological context. However, parallels exist with *Ophiocordyceps*-infected ants that exhibit changes in behavior that include increased walking, deviation from foraging trails, reduced nestmate communication, and the final manipulated bite ^12–16^.

#### Changes in dopamine metabolism possibly modifies ant behavior

A combination of metabolomic and transcriptomic data indirectly suggest increased dopamine metabolism in manipulated ants. The metabolomic signals consisted of changes in neurotransmitter precursor abundances and enrichment of pathways in metabolite network modules correlated to manipulation. Phenylalanine and tyrosine are precursors in synthesis pathways for dopamine and octopamine. KEGG pathway “phenylalanine metabolism” was enriched in both network modules A1 and P2 (p = 0.026 and 0.028, fold-enrichment = 1.5 and 1.9, respectively), both of which were negatively correlated to manipulated ant samples. Additionally, KEGG pathway “tyrosine metabolism” was enriched in modules P2 and P3 (p = 0.030 and 0.024, fold-enrichment = 1.4 and 1.6, respectively), both also negatively correlated to manipulation (Figure 3, Table 1). Moreover, module P2 also contained the most informative compound annotations regarding dopamine metabolism.

The immediate precursor to dopamine, L-DOPA, was 2.8-fold reduced in abundance in manipulated ants (p = 0.008, top feature) (Fig. 4). L-tyrosine was also annotated in module P2 and is the precursor to L-DOPA. L-Tyrosine was a DAM with a 2.1-fold decrease in abundance within the polyphenols dataset (p = 1E-4, top feature). An annotation for L-tyrosine was also present in the network module A1, however, we did not identify this metabolite feature as a DAM in the biogenic amines dataset (p = 0.154, 1.2-fold reduction). Outside of network modules A1, P2, and P3, we also detected phenylalanine, the precursor to tyrosine. Phenylalanine showed a modest increase in abundance during manipulation just below our 1.5-fold change threshold for classification as a DAM (p = 0.019, 1.4-fold increase) (Fig. 4). Decreases in L-tyrosine and L-DOPA are difficult to interpret, but possibly indicate their conversion to produce high concentrations of dopamine. Alternatively, reduced precursor abundances could reflect wide-spread reduced levels of metabolites, including dopamine. However, our previous RNAseq data suggested an increased dopamine synthesis activity causing the depletion of these dopamine precursors.

**Figure 4.**
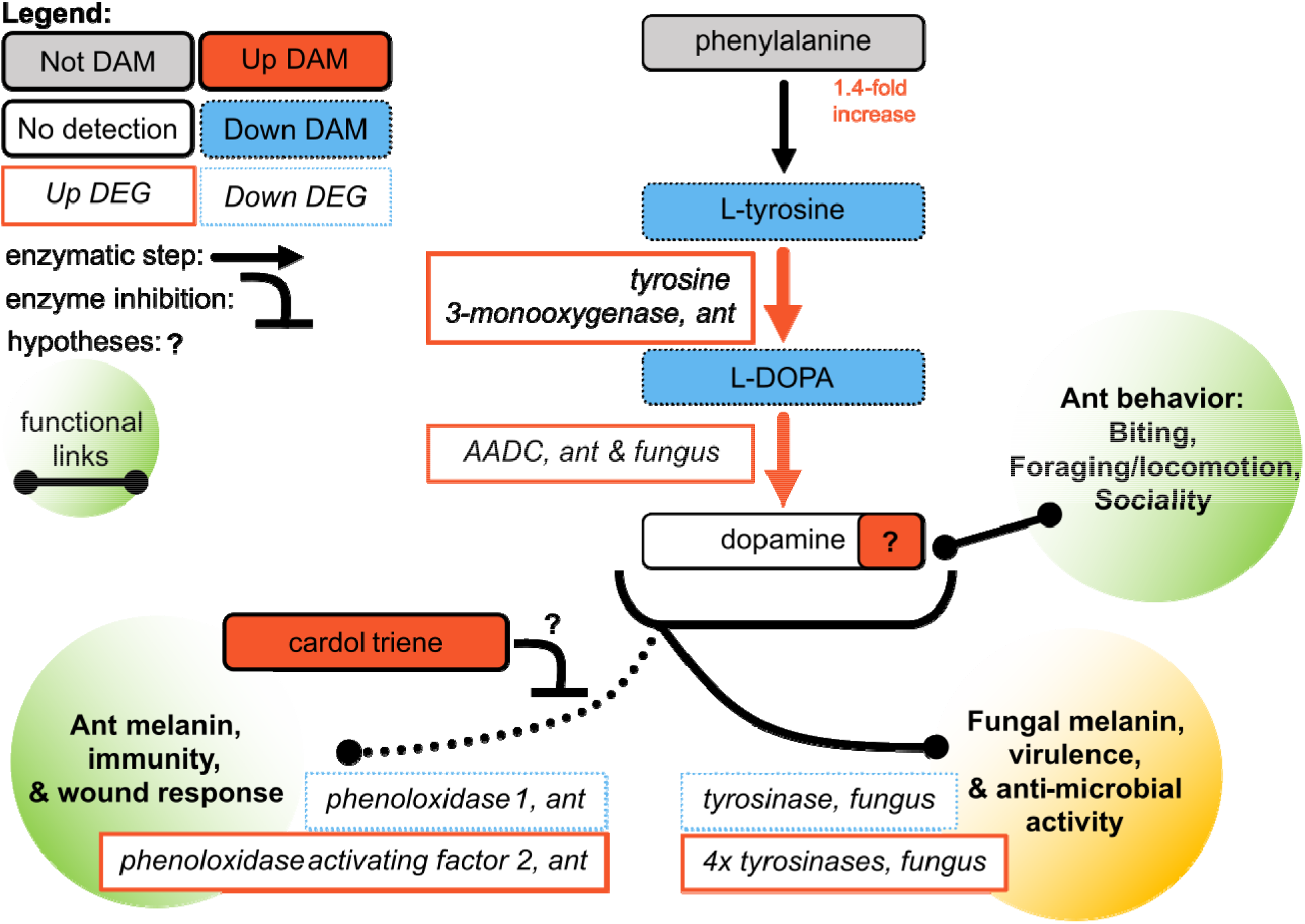
Altered neurotransmitter precursor metabolism suggests suppressed ant immunity and increased dopamine during infection and manipulation. Upregulation of enzyme genes converting tyrosine to L-DOPA to dopamine coupled with reduced tyrosine and L-DOPA levels suggest increased concentrations of dopamine during infection, which may, in turn, contribute to manipulated behavior. Additionally, cardol triene may be produced by the fungus to suppress ant immunity by inhibiting melanin synthesis and thereby also increase availability of tyrosine and L-DOPA for dopamine synthesis. Combined with these specific metabolites, phenylalanine and tyrosine metabolism pathway enrichments further implicate these processes in manipulated behavior. Increases in expression, abundance, or activity are indicated by solid lines and red color, decreases by dotted lines and blue color. Metabolites are in rounded rectangles, genes are in rectangles. Annotated metabolites that were not DAMs have gray color, while compounds with white color were not identified in the LC-MS/MS data. Hypothesized links to ant (green) and fungal (orange) phenotypes are given in the shaded circles. Black arrows are shown when useful for context to indicate that an enzymatic step had a corresponding enzyme gene in either the fungal or ant genome, but it was not a DEG.

During manipulation, both ant hosts and fungal parasites upregulated an aromatic L-amino acid decarboxylase (AADC) gene (Fig. 4). This enzyme catalyzes the final step of L-DOPA conversion to dopamine ^60^. The AADC enzyme requires vitamin B6 as a cofactor. We detected differential abundance of an inactive vitamer form of B6 and a precursor for its *de novo* synthesis in module A5 (Supplementary Discussion S4). However, the combination of our metabolomic and transcriptomic data do not provide for a clear interpretation as to if active vitamin B6 is changing in abundance and in which organism(s).

In addition, a tyrosine 3-monooxygenase, which catalyzes the rate-limiting conversion of L-tyrosine to L-DOPA, was upregulated by manipulated ants. Meanwhile, the ant homolog to Henna (i.e., DTPHu phenylalanine 4-monooxygenase), which converts phenylalanine to L-tyrosine was not differentially expressed. This enzyme also converts peripheral tryptophan to serotonin’s precursor in *D. melanogaster* ^61–63^. This consistent expression may help explain the only modest change of phenylalanine during manipulation while its derivatives were found at lower concentrations.

#### Shifts in dopamine and melanin metabolism could mediate host-pathogen immunological interactions during manipulation

Dopamine and its precursors can also be converted into melanin, a key compound for insect wound response and immunity ^52, 64, 65^. Additionally, fungi can synthesize melanin for use in host-pathogen interactions and against competing microbes ^66, 67^. We found evidence that this pathway for melanin production is suppressed in the ant during manipulation and, possibly, increased in the fungus (Fig. 4).

We propose that the production of a fungal-derived compound may inhibit host tyrosinases necessary to convert dopamine-pathway metabolites to melanin and hence simultaneously steer these metabolites toward dopamine production. This metabolite feature was annotated as cardol triene (i.e., 5- (8,11,14-pentadecatrienyl)resorcinol) and was a DAM only detected in manipulated ant samples (p = 0.0073, median peak value = 19,604, top feature) (Fig. 4). As cardol triene was never detected in healthy ants, it plausibly is produced solely by the fungus, although we cannot discount the possibility that production of this metabolite is produced by ants combatting infection and completely absent in healthy individuals. We detected the putative cardol triene in six of 10 manipulation samples, which could indicate that cardol triene levels are tightly regulated. Cardol triene has been shown to inhibit tyrosinase oxidation of L-DOPA in vitro ^68^. In insects, phenoloxidases are a type of tyrosinase that are key enzymes that begin the conversion of dopamine and L-DOPA to pro-immune melanin production pathways ^52, 65, 69^. As such, we hypothesize that fungal cardol triene chemically inhibits host phenoloxidase activity to reduce host melanin metabolism.

Additionally, ants showed mixed transcriptional signals related to the melanin-immune response by phenoloxidase and phenoloxidase activating factor genes (Fig. 4) ^70^. Entomopathogen infections inhibit phenoloxidase activity in mosquitoes and suppression of the activation of pro-phenoloxidase has been hypothesized as a mechanism of fungal interference with insect host immunity ^71, 72^. In addition to suppressing host melanin production, the fungus may be converting L-DOPA or related compounds into fungal melanin precursors, as four upregulated and one downregulated tyrosinases were found during manipulation (Fig. 4). Three of the upregulated fungal tyrosinases are putatively secreted and may possibly interact with host tyrosine and its derivatives. Taken together, our metabolomics and previous transcriptomics data not only suggest that altered dopamine metabolism could lead to increased dopamine concentration that induces altered host behavior, but also result in weakened host immunity while bolstering parasite pathogenicity (Fig. 4).

#### Serotonin and octopamine metabolism may also correlate to manipulated ant behavior

Biosynthetically and phenotypically intertwined with dopamine, fluctuating serotonin and octopamine concentrations may also contribute to manipulated ant behavior. Although the metabolomic signals for these compounds were less robust than for dopamine, we found multiomic evidence that offers support for hypotheses of increasing serotonin and decreasing octopamine (Supplementary Discussion S5). For example, higher serotonin concentration is indicated by upregulation of the AADC enzyme shared between dopamine and serotonin pathways ^60^ and at least one signal for the reduction of tryptophan, which can be converted into the precursor of serotonin. Coupled with the increasing serotonin, we additionally propose a hypothesis for increased kynurenic acid, which is also tied to tryptophan and may act as a neuroprotectant for heavily diseased and manipulated ants (Supplementary Discussion S5). Reduced octopamine is suggested by changes in tyramine metabolism evidenced by both reduced L-tyrosine (a tyramine precursor) and a tyramine derivative (N-acetyltyramine) (Supplementary Discussion S5).

### Altered glycerophospholipid metabolism suggests increased fungal cell membrane activity and ant accumulation of the excitatory neurotransmitter, acetylcholine

The KEGG pathway “glycerophospholipid metabolism” was enriched in module P2 and among the top-features (p =0.024 and 0.030, fold-enrichment = 1.9 and 1.4, respectively). Of our annotated features, metabolic signals centered on choline offered the most insight into altered glycerophospholipid metabolism. We found evidence for two possible major effects, changes in cell membrane composition and the accumulation of acetylcholine. Glycerophospholipids and choline-containing compounds have been associated with neurodegenerative diseases, such as Alzheimer’s or Parkinson’s, as well as insect host-pathogen responses ^73–75^. Choline is a key component involved in the synthesis of phospholipids associated with forming cell membranes, especially as incorporated through phosphatidylcholines ^76^.

Although, in fungi and insects, phosphatidylethanolamines are also a major cell membrane component ^77–79^. Using the lipids LC-MS/MS dataset we compared peak values of phosphatidylethanolamines (n = 95 features) and phosphatidylcholines (n = 85 features) and found phosphatidylethanolamines to indeed be abundant in our insect and fungal tissue samples. Within healthy ants, phosphatidylethanolamines were only slightly more abundant (1.10-fold greater) than phosphatidylcholines (t-test, p = 0.017, t = −2.657). Within fungus-manipulated ants, there was no strong difference (t-test, p = 0.739, t = −0.339, phosphatidylethanolamines 1.05-fold higher).

Choline was among the DAMs that increased in abundance during manipulation (p = 1.7E-11, 20.0-fold increase, top feature) (Fig. 5). We also found other DAMs related to choline within the glycerophospholipid metabolism pathway. Glycerophosphocholine, a choline precursor, increased at manipulation in both the amine and polyphenol datasets (p = 3.4E-11 and 1.2E-7, 5.9- and 7.7-fold increase, respectively, top features). We also detected two DAMs interconnected between choline and phosphatidylethanolamines: ethanolamine (p = 5.2E-5, 2.1-fold increase, top feature) and ethanolamine phosphate (p =1.6E-9, 5.5-fold decrease, top feature) (Fig. 5).

**Figure 5.**
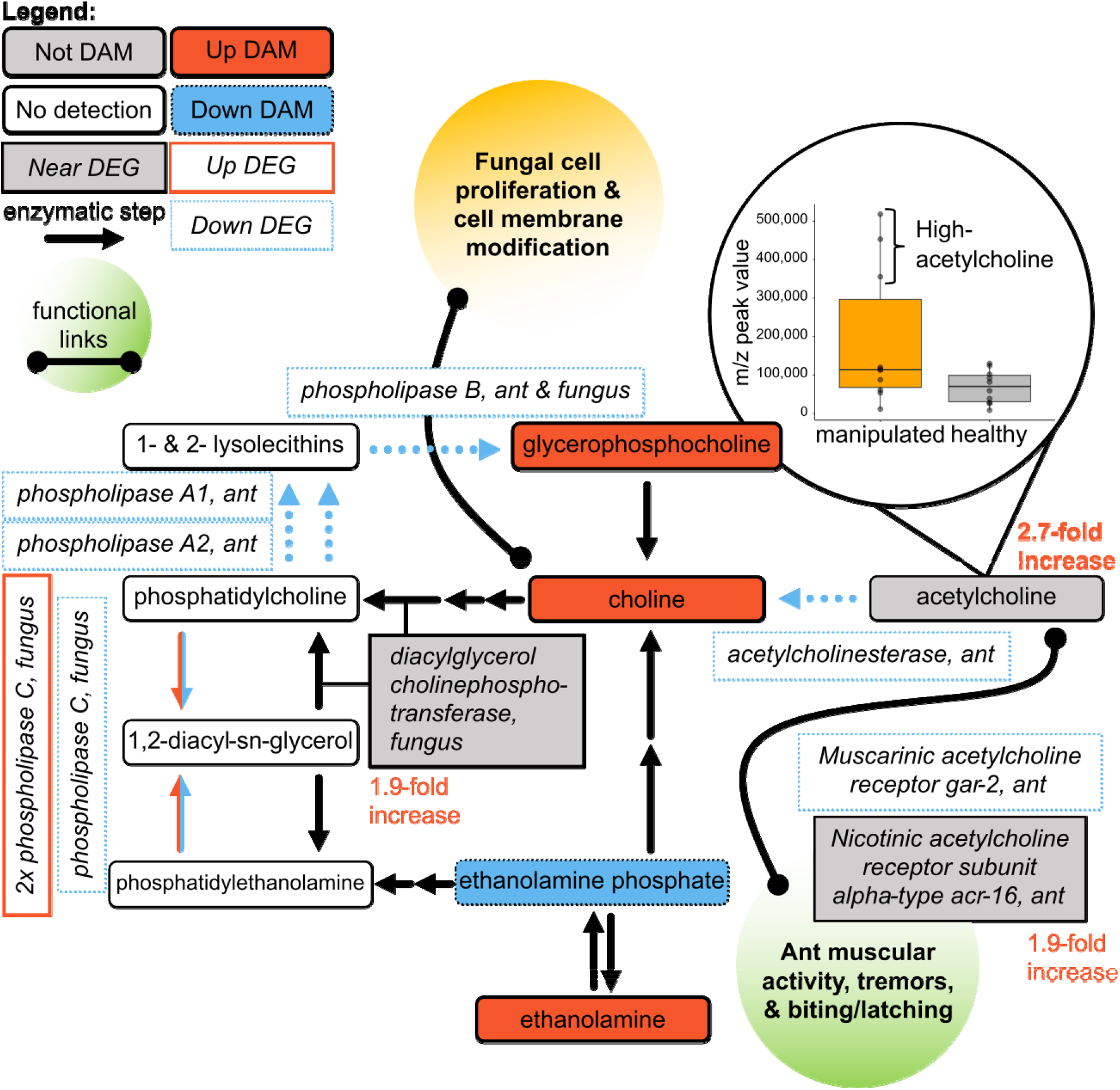
Changing glycerophospholipid metabolism during manipulation primarily indicate altered acetylcholine levels in the ant and cell membrane metabolism in the fungus. Manipulated ant samples appeared to have two distinct groups, one with high acetylcholine and one with levels similar to healthy ants. In manipulated ants, the putative acetylcholinesterase and two acetylcholine receptor genes were downregulated, suggesting acetylcholine levels were accumulating. We hypothesize that increased acetylcholine in manipulated ants is associated with the increased muscular activity during early manipulation (e.g., tremors, biting, and latching behavior). Once the ant is secured in a summited position and the mandible muscles (and associated neurons) are further degraded, the fungus may be scavenging and converting the available acetylcholine to choline, thereby rapidly reducing acetylcholine levels. The fungus may be converting choline to phospholipids, such as phosphatidylcholines to remodel and build cell membranes. Although we detected multiple phosphatidylcholine and phosphatidylethanolamine class features, we did not identify the singular metabolites 1,2- diacyl-sn-glycero-3-phosphocholine and 1-acyl-2-acyl-sn-glycero-3-phosphoethanolamine, annotated as “phosphatidylcholine” and “phosphatidylethanolamine” in the KEGG glycerophospholipid pathway, respectively. The legend is the same as described in Figure 5, with the additional inclusion of genes that nearly qualified as DEGs depicted in gray.

Multiple phospholipase DEGS in both host and parasite indicate possible altered lipid metabolism acting on molecules biosynthetically related to choline and ethanolamine compounds. Broadly, phospholipases cleave phospholipids and have been implicated in cell signaling, fungal pathogenicity, and cell membrane metabolism ^80, 81^. Although Menezes *et al.* 2023 note a preponderance phospholipase A2 in the *Ophiocordyceps australis* genome, with possible functions in pathogenesis – our previous transcriptomics data did not detect fungal type A2 DEGs during active manipulation. Rather, three putative *O*. *camponoti-floridani* phospholipase C genes were DEGs, two upregulated and one downregulated (Fig. 5). When cleaving either phosphatidylcholines or phosphatidylethanolamines they can produce an intermediate 1,2-diacyl-sn-glycerol product. Additionally, the fungus expressed a putative diacylglycerol cholinephosphotransferase gene 1.9-fold higher during manipulation, nearly qualifying as a DEG. This enzyme combines 1,2-diacyl-sn-glycerol and CDP-choline to produce phosphatidylcholine.

Phosphatidylcholines could then be converted to glycerophosphocholine within two metabolic steps, but this pathway appeared to be downregulated. Three genes putatively encoding the enzymes driving this conversion were downregulated in the ant (phospholipases A1, A2, and B). In the fungus, a phospholipase B-like gene was also downregulated (Fig. 5). The net effect appears to be fungal composition and/or catabolism of phosphatidylethanolamines and synthesis of phosphatidylcholines (Fig. 5). Fungal phospholipase activity may also disrupt host responses and facilitate destruction of host tissues^81^. Modification of the fungal cell membrane could also be necessary to accommodate transmembrane proteins or secretory activity involving organelle membranes such as in the endoplasmic reticulum ^83, 84^. This altered membrane metabolism may be involved in pathogenesis and release of manipulation effectors, but alternatively or additionally, relate to changing fungal growth ^85, 86^. Shortly after manipulating its host, the fungus undergoes a switch from blastospore to hyphal growth and further colonizes the host cadaver ^12, 15, 20, 26^. In line with this, we also hypothesize that signals related to gamma-aminobutyric acid (GABA) and glutamate indicate altered fungal growth metabolism (see below, Supplementary Discussion S6).

Choline can also be metabolized into acetylcholine, the most abundant excitatory neurotransmitter in insects ^87^. In bees, excessive activation of acetylcholine receptors can cause changes in locomotor, navigational, foraging, and social behaviors ^88^. Acetylcholine can be rapidly converted back to choline by the activity of the acetylcholinesterase enzyme, which has been a target of enzyme-inhibiting insecticides ^89–92^. From this fast enzymatic activity, we intuit that acetylcholine levels could reflect changes in physiology and behavior at a fine timescale since a subgroup of the manipulated ant samples showed markedly higher acetylcholine levels than both healthy and other manipulated ants (Fig. 5, Supplementary Fig. 1). Speculatively, this could indicate that, for some fast-changing metabolites such as acetylcholine, our sampling approach captured multiple subtly different physiological states during the dynamic process of behavioral manipulation.

Acetylcholine showed a 2.7-fold mean increase at manipulation overall. However, it was not identified as a DAM (p = 0.142) with the feature-peak value distribution showing two distinct groups. Three manipulated ants had 5.5-fold higher acetylcholine peak values than the other seven that spanned a similar range as control samples (Fig. 5). The previously collected transcriptomics data similarly showed a broad distribution of acetylcholinesterase-2 expression in manipulated ants. Notably, acetylcholinesterase-2 is the functionally dominant acetylcholinesterase enzyme in *Camponotus* and many hymenopterans ^93^. In the ant, an acetylcholinesterase-2-like gene was downregulated during active manipulation overall, but two of the five samples had 2.5-fold lower acetylcholinesterase expression than the other three (Supplementary Fig. S1) ^24^. As the ant dies, acetylcholinesterase-2 expression begins to rebound toward healthy ant levels (Supplementary Fig. S1). Additionally, the ant downregulated a putative muscarinic acetylcholine receptor gar 2 that has been found in sensory and motor neurons of *C. elegans* ^94^. Nearly a DEG as well, a putative ant nicotinic acetylcholine receptor acr-16 subunit gene was expressed 1.9-fold lower during manipulation. Possibly, receptor downregulation represents a homeostatic response to the proposed elevated acetylcholine levels present during part of manipulated summiting and biting.

Although acetylcholine has fast-acting regulatory effects on motor behavior, importantly, glutamate is proposed as the direct primary excitatory neurotransmitter at insect neuromuscular junctions ^87, 95, 96^. Glutamate was nearly classified as a reduced DAM (p = 0.007, 1.4-fold decrease) while a related compound, glutamine, did qualify (p = 1.8E-6 and 6.2E-11, 3.3- and 3.8-fold decrease, in the amine and polyphenol datasets, respectively), with transcriptomic evidence suggesting that the host and parasite may be mounting contrasting regulatory responses ^24^ (Supplementary Discussion S6).

### Common metabolites between *Ophiocordyceps*-*Camponotus* manipulations across species- interactions

Previous metabolomic studies with *O. kimflemingiae* have identified many metabolites associated with infection of *C. castaneus* and proposed links between these compounds and behavioral manipulation ^21, 22, 25^. Laboratory co-culture of *O. kimflemingiae* with brains from *C. castaneus* and other ant species showed both shared metabolites and distinct compounds reflecting a combination of target host-specific fungal secretions and ant species profile ^25^. Other studies have compared the metabolomes of *O. kimflemingiae* and generalist entomopathogenic *Cordyceps bassiana* infections, again finding both overlap and differences in metabolomic profiles ^21, 22^.

These studies detected ergothioneine, an amino acid produced by some fungi, but not animals, in mixed ant-fungal tissue from manipulated ant brains and muscles ^21, 22^. Corroborating these findings, we detected two ergothioneine signals (biogenic amine and polyphenol datasets) that were wholly absent or with peak values near that of blank-collection samples in the healthy ants (p = 7.3E-15 and 4.6E-12, top features). In addition to fungal metabolism, ergothioneine has been discussed as a possible neuroprotectant to preserve the ant brain until manipulated summiting and biting are complete ^21^. Supporting this line of reasoning, ergothioneine in the ant brain was not detected at elevated levels in moribund ants infected by the generalist entomopathogen *C. bassiana* that consumes hosts more quickly, without inducing summit disease ^21^. Notably, *Ophiocordyceps* do not appear to typically reside directly within the brain, suggesting ergothioneine may be secreted by the fungus ^13, 21, 26, 97^ ^preprint^.

Previous work with *O. kimflemingiae* and this work with *O. camponoti-floridani* also found adenosine and adenosine-phosphate (AMP) to be differentially abundant between manipulation and healthy and/or generalist-infected ants ^21, 22^. We found two features annotated as adenosine, one which was not differentially abundant and one that was a reduced DAM in manipulated samples (p = 6.6E-4, 2.5-fold decrease). In contrast, previous works found adenosine to increase in abundance in both manipulated ant brains and mandible muscles ^21, 22^. Whether these contrasting results indicate species-specific patterns or high variation is unclear. We also detected decreased AMP (p = 0.003, 2.1-decrease, top feature), as was found in manipulated *C. castaneus* ^21, 22^. In *O. kimflemingiae* infection, a role for adenosine-mediated neuromodulation was hypothesized, as was changing AMP levels possibly reflecting changes in mitochondrial activity ^21, 22^. Cyclic AMP is involved in many pathways, including as a secondary messenger in G-protein coupled receptor signaling that has been hypothesized to change during *O. camponoti-floridani* behavioral manipulation with the previous transcriptomic study ^24^. If changing abundance of AMP reflects upon the cyclic form as well, this could relate to dysregulation of receptor signaling. Metabolically related to adenosine, hypoxanthine and inosine were previously proposed to have links to energetic and neurological processes in manipulated ants ^21, 22^. As in those studies, we also detected increased levels in two of three hypoxanthine-annotated features at manipulation (p = 0.007 and 7.7E-6, 1.6-fold increase or absent in sham, the latter of which was a top feature). We also detected one inosine and two inosine-monophosphate features increased in manipulated ants (p = 0.002, 3.8E-7, and 1.4E-5, 1.9-, 2.8-, 2.5-fold increase, top features). This could indicate the conversion of AMP and adenosine to hypoxanthine primarily via inosine intermediates.

## Conclusion

Infection and behavioral manipulation of hosts could involve many interacting molecular and cellular processes. As these host-parasite interactions are better characterized, we can also gain insights into specialist co-evolutions, the mechanistic basis of animal behavior, and the discovery of new bioactive compounds. To unravel the mechanistic complexity of infection and manipulated host behavior, we require functional tests built upon robust hypotheses. Towards this goal, we studied parasitic manipulation of *C. floridanus* by *O. camponoti-floridani*, combining metabolomic, transcriptomic, and genomic data.

We found evidence for altered biogenic monoamine neurotransmitter production during infection and manipulation that may be proximate causes of manipulated behavior. Although, we did not identify and measure these neurotransmitters directly. Going forward, now with evidence implicating these neurotransmitters in behavioral manipulation, experiments focused on positively identifying them could employ increased sample concentration (i.e, pooling multiple ant heads per sample) and recent advancements in LC-MS/MS approaches to detect monoamines in insect brain tissue ^98^. Additionally, different metabolomic analyses may support discovery of specialized fungal effector metabolites as has been observed in a fly-manipulating fungus and hypothesized in *Ophiocordyceps* ^99,100^ ^preprint^.

However, by integrating transcriptomic data with the metabolite features we did identify, we propose a scenario where the manipulated ant has elevated production of dopamine, with possible increases of serotonin and decreases of octopamine. The fungus may also simultaneously suppress the host’s melanin and immunity pathways while metabolizing dopamine and its precursors for fungal melanin production. Ants infected by *Ophiocordyceps* display enhanced locomotion/activity levels, reduction of social behaviors, and a final summit latching behavior ^12–16^. Elements of these behaviors have already been associated with neurotransmitter signaling in ants, albeit phenotypic variation across species and social or physiological context exists ^54–59^. Intriguingly, aquatic gammarid crustaceans manipulated by acanthocephalan worms display a phototactic summiting and clinging phenotype in the water column associated with increased serotonin, decreased octopamine, and suppressed phenoloxidase-mediated immunity ^101–104^. In essence, our data indicate comparable biogenic amine level changes and immunity suppression in manipulated *Camponotus,* which is in line with the premise that convergently evolved manipulated summiting behaviors are established by overlapping mechanisms ^5,105^. Such mechanisms are likely specific to different behavioral manipulation phenotypes, but may include common players found across phenotypes. For example, some parasitoid wasps induce a low-activity, defensive “bodyguard” state in their insect hosts that has been hypothesized to be linked to increased, rather than decreased, octopamine levels in the host ^106^. Manipulators of vertebrate behavior are also thought to leverage serotoninergic and dopaminergic processes to alter fish and mouse behaviors related to escape responses, locomotion, and aggression ^50, 107–109^. How much of the alteration of neurotransmitter metabolism may be a general response to fungal infection and how much results from specific manipulation by *Ophiocordyceps* is difficult to pinpoint at this time. Possibly promoting both infection and manipulation, cardol triene appears to be a fungal derived compound that could suppress ant immunity and increase substrate availability for dopamine production. Also, *Ophiocordyceps* upregulated three putatively secreted tyrosinases that possibly interact with host dopamine/melanin precursors. Although less typically reported than their intracellular counterparts, extracellular fungal tyrosinases have also been found in another fungus ^110, 111^.

Changing glycerophospholipid metabolism during manipulation, largely centered on choline, suggested fungal biosynthesis of cell membrane components and dysregulation of acetylcholine levels in the ant. *Ophiocordyceps* may require increased choline-derived cell-membrane components used during cell proliferation and membrane remodeling for structures such as lipid rafts or transmembrane proteins important during infection, manipulation, and hyphal colonization of the host ^83–86^. In some manipulated ant samples, we detected elevated acetylcholine concentrations, which may correspond to reduced acetylcholinesterase gene expression previously observed. Speculatively, transient high acetylcholine levels could contribute to summiting and wandering behavior (i.e., increased locomotion), tremors, and the hypercontracted biting that locks manipulated ants into place ^12, 15–17, 20, 25, 87, 88, 112^.

By inducing changes in neurotransmitters that regulate behavior in healthy ants, the fungus may be impinging upon existing host systems to modify behavior rather than forcibly introducing novel pathways ^8, 18^. Indeed, many of the hallmark phenotypes of infection by ant-manipulating *Ophiocordyceps* are not so dissimilar from activities influenced by levels of dopamine, serotonin, octopamine, and acetylcholine in eusocial insects: foraging/locomotor behavior (i.e., summiting and hyperactivity), sociality (i.e., wandering and reduced communication), and the biting/aggression (the final “death grip” bite) ^47, 48, 49–51, 54–58, 59^. Changes in social behavior could not only be a symptom of disease, but contribute to it as well. As socially isolated infected ants may less frequently exchange trophallactic fluid with nestmates, these infected individuals would lose an important source of nutrition, communication, and hormones ^113^ that may alter their metabolome. How exactly the fungus dysregulates physiological pathways that influence behavior, such as insect hormone metabolism, and to what degree some level of dysregulation is typical in many diseases is still unclear. These hypotheses can be further investigated by functional tests (e.g., gene knockouts or dosing ants with small compounds) and LC-MS/MS validations of key compounds using chemical standards and improved protocols.

## Methods

### Ant collection & husbandry

We used a single laboratory-housed wild-caught colony of *C. floridanus* to produce all samples. This colony was collected in March 2021 in the University of Central Florida arboretum and had several hundred individuals including minors, majors, and brood. Based on methods in Will *et al.* 2020, we housed the colony in a 9.5 L plastic container (42 cm long × 29 cm wide) lined with talcum powder (Fisher Scientific) and contained aluminum foil wrapped test-tubes (16 mm x 125 mm, Fisher Scientific) with moist cotton to serve as darkened, humid nest spaces. Ants fed ad libitum on 15% sucrose solution, water, and sporadic frozen *Drosophila* feedings. Following collection, we housed the ants indoors under 48 h constant light to prepare them for circadian entrainment to laboratory conditions. Follow the “circadian reset,” we transferred the colony to a climate-controlled incubator (I36VL, Percival) cycling 12-12 h, light-dark, 28-20 C°, 65-80% RH daily cycles. Climate conditions cycled in a ramping manner, with a 4 h ramp from lights-on to a 4 h hold at peak day conditions to a 4 h ramp to lights-off. The colony housed under these conditions for approximately six weeks before individuals were removed for the experiment, which was conducted in the same incubator.

### Fungal strain collection and culturing

*O. camponoti-floridani* strain OJ2.1 was used to infect all ants in this work. We isolated this fungal strain using mycelium extracted from the abdomen of a recently manipulated and expired *C. floridanus* cadaver. This cadaver was found in the Ocala National Forest, FL (29° 13’ 41.7” N, 81° 41’ 44.9” W), in March 2021 and permitted by letter of authorization (United States Department of Agriculture, #2720-22). The isolation procedure, short ribosomal subunit genetic validation, and culturing followed the protocols detailed in Will *et al.* 2020. In preparation for experimental infections, we grew fungal cells for 10 days in T25 cell culture flasks with 20 mL Grace’s medium (Gibco) supplemented with 2.5% fetal bovine serum (Gibco) (“GF” medium), shaken at 50 rpm in 28 C°. We then filtered the culture using two-layers of 12-ply gauze (Curity) to separate yeast-like blastospores from hyphal cells. We adjusted the filtered blastospores to a working concentration of approximately 3 x 10^6^ cells/mL (optical density 660 nm = ca. 0.5) prior to injection.

### Experimental ant infections

Prior to *Ophiocordyceps* infection and sham injection, we separated ants from their colony into treatment groups. For the infection treatment, we gathered 51 ants, of which we injected 41 individuals. For the control group, which received sham injections, we collected 38 ants, of which we sham-injected 30 individuals. The additional ants in each group did not receive treatment other than a painted dot on their abdomens (Testors paint) to be able to recognize them. Our previous experiments have shown that such a small group of untreated ants improves the recovery of the treated group during the first few days of the experiment ^14^. One day before infection treatments, we removed the sugar and water from the ant box to reduce the amount of fluid in the ants and ease the injection procedure. We injected each ant with 0.5 µL *Ophiocordyceps* blastospores (infection, n = 41) or 0.5 µL of GF (sham, n = 30) as detailed in Will *et al.* 2020. We placed the treated ants and their painted “caretaker ants” in plaster-lined boxes with biting structures (*Tillandsia usneoides* tied to wooden sticks with twine) to be able to observe *Ophiocordyceps*-induced summiting behavior.

### Behavioral observations

After three dpi we removed ants that did not survive the treatment. Starting day four, we screened for manipulated ants and deaths twice per day, at zeitgeber time (ZT) 0, the beginning of subjective day (i.e., lights-on), and ZT 6, the mid-point of subjective day. At ten dpi, we removed the painted caretaker ants as previous work has observed *C. floridanus* to aggressively interfere with manipulated nestmates, presumably as a component of their social immunity and nest hygiene ^14, 24, 114^. Starting 15 dpi, we began more frequent opportunistic checks as behavioral manipulations of *C. floridanus* that involve summiting had previously been observed starting 19 dpi in the laboratory ^24^. After discovering the first manipulation at 21 dpi, we checked multiple times, daily beginning at ZT 20 and ending at ZT 14 (on the next subjective day). We selected this observation window to improve the odds of collecting manipulations close to their onset since our previous laboratory infections with *O. camponoti-floridani* showed that manipulated summiting synchronized to the ants’ subjective pre-dawn hours ^24^. Summiting behavior is characterized by infected individuals clinging and biting *Tillandsia* suspended in the experiment box. The behavior has never been observed in sham-treated individuals. We screened non-manipulated mortalities for fungal infection by crushing tissue from the dead ant in 50 μL of water, briefly centrifuging, and examining 15 μL of the supernatant under a microscope. Notable levels of fungal cells were always and only seen in the *Ophiocordyceps* treatment ants. We analyzed and plotted survival data using the R packages survival and survminer in R (v 4.1) with R Studio (v. 2021.09.2) ^115–118^.

### Sample collection for LC-MS/MS

We collected live manipulated ants displaying characteristic summiting biting behavior for LC-MS/MS immediately upon discovery (n = 20). We subsequently collected dpi-matched sham ants at ZT-21.5, as this was the most common time at which manipulated ants were collected (n = 20) (see Results and Discussion). Manipulated ants found after death were recorded for survival statistics but not used in the LC-MS/MS analyses (n = 3). For manipulated ant collection, we held ants with forceps, clipped the biting substrate just above and below the manipulated ant and submerged the ant in liquid nitrogen briefly until frozen. The sample was then placed in a pre-chilled, sterile 2 mL microcentrifuge tube (USA Scientific) kept in liquid nitrogen until long-term storage at −80 C°. We collected sham ant samples in a similar manner, lifting them from the experiment box with forceps.

We also collected three blank samples at the beginning, middle, and end of manipulated ant sampling, i.e., after the first, 10^th^, and 20^th^ manipulated ants were found (n = 3). We made blanks by briefly touching the forceps to the floor of the sham treatment housing boxes, submerging them in liquid nitrogen, and swirling them in a microcentrifuge tube. We cleaned all tools and surfaces with 70% ethanol and deionized water before sampling and between sample types.

We later prepared collected samples for LC-MS/MS by isolating ant heads. Each frozen ant was quickly processed on a plastic petri dish (USA Scientific) on a bed of dry ice. Using forceps, we removed biting substrate from the ant’s mandibles if needed, snapped the head off, and placed it in a new microcentrifuge tube. For blank samples, we dragged the forceps along the petri dish and swirled the forceps inside the original collection tube. We cleaned the tools and petri dish with 70% ethanol and deionized water between individuals and used a new petri dish between sample types. All samples and tubes were kept constantly chilled on dry ice or in liquid nitrogen.

### LC-MS/MS processing of ant head samples

All samples were sent on dry ice to the West Coast Metabolomics Center (WCMC, University of California, Davis) for LC-MS/MS analysis. The WCMC processed samples in three LC-MS/MS protocols broadly tailored toward different chemistries: “biogenic amines,” “polyphenols and flavonoids,” and “complex lipids.” From the 20 samples collected per infection and sham treatments, the WCMC collected polyphenol data for ten individual ant heads. They used the remaining ten heads per treatment for biphasic extractions, with each individual head used for both biogenic amines and lipids. Blank-collection samples were shared across the three LC-MS/MS chemistries. The WCMC produced quality-control (QC) pool samples consisting of all samples used per protocol and additional blank-extraction controls (and therefore distinct from our experimental blanks made during sample collection). These WCMC control samples were run at the beginning and every ten samples thereafter per protocol. All LC-MS/MS protocols collected MS/MS data in a top-four data-dependent-acquisition (DDA) mode in both electrospray ionization (ESI) positive and negative modes.

The WCMC processed samples for biogenic amines for use in hydrophilic interaction chromatography (HILIC) with quadrapole time of flight (Q-TOF) mass spectrometry. Homogenized ant head samples were extracted with a methanol and methyl-tert-butyl ether protocol ^119^. The aqueous phase was used for biogenic amines LC-MS/MS and the organic phase for lipids (see below). Extractions were mixed with internal standards for normalization according to WCMC protocols to a final volume of 100 μL (Supplementary Data S3).

Samples for lipids were processed for use in charged surface hybrid (CSH) chromatography with Q-TOF mass spectrometry. The organic phase per biphasic extraction (shared with biogenic amines, see above) was taken for lipids LC-MS/MS and mixed with internal standards according to WCMC protocols to a final resuspension volume of 110 μL (Supplementary Data S3).

Samples for polyphenols were processed for use in pentafluorophenyl (PFP) chromatography with QExactive HF (QE) mass spectrometry. Homogenized ant head samples were extracted in a similar fashion as described above and resuspended in a final 100 μL volume with internal standards (Supplementary Data S3). Further details for all three LC-MS/MS protocols are provided in Supplementary Methods.

### Metabolomic data processing & feature filtering

The WCMC performed initial processing steps on all samples to produce metabolomic feature lists consisting of mass/charge ratio (*m/z*), retention time (rt), and value of quantification ion peaks. They aligned raw data with MS-DIAL (v 4.7.0) ^120^, subtracted signals detected in the blank-extractions produced in the core facility, removed duplicate signals and isotopes, normalized samples by internal standards, and assigned compound and adduct annotations based on manual MS/MS matching in Massbank, Metlin, and NIST14 databases and, for lipids, with LipidBlast (v 68) ^121^. The resulting data consisted of several thousand features per dataset: biogenic amines (5605 total with 141 annotated), polyphenols (6178 total with 158 annotated), and lipids (7021 with 517 annotated) (Supplementary Data S4).

We applied additional feature quality control and data reduction methods to create a final list of metabolite features to investigate. Features that had greater than 50% missing values within both sham and infection treatments were removed. Similarly, we removed features with both median peak values (sham and infection) less than double that of median blank sample peaks. Following removal based on missingness and peak values, we removed features that we determined to have signals too variable for robust analysis based on their coefficient of variation (CV%, or relative standard deviation). For the biogenic amines and lipid datasets that had successful QC pool data returned, we applied a CV% maximum threshold to remove peaks that varied greatly due to technical variation between QC pool injections. We selected the CV% threshold as the lowest (i.e., strictest) value between 30% and a dataset specific value determined from the distribution of CV% values. We calculated this data-specific value as the 75^th^-quartile + 1.5*interquartile-range (IQR), as is used to identify extreme values or outliers in data distributions (e.g., boxplots). For the lipids, the 1.5*IQR method produced CV 25% as the cutoff, removing 5.7% of features. For biogenic amines, the 1.5*IQR CV% was greater than 30%; therefore, 30% was used, removing 6.6% of features. The WCMC reported that all polyphenol QC pools failed on acquisition. Therefore, we ranked each feature by its lowest CV% (i.e., from either CV%-sham or CV%- manipulated) and removed the highest 6.1% of features. We selected this 6.1% value because it was the mean of the CV% removal rates of the biogenic amine and lipid datasets. Although we cannot fully discount biological variation in biogenic amine peak values within a treatment, by selecting the lowest CV% per feature (whether from the sham or infection treatment), we sought to remove only features that had high CV% regardless of treatment. After these filtering steps, we continued with 3268 biogenic amine features (58% of initially collected), 4878 polyphenol features (79% of initial), and 6305 lipid features (90% of initial).

We further applied a putative adduct calling step to consolidate unannotated features that may represent different adducts of the same compound. Per LC-MS/MS dataset, we divided features by treatment and ESI mode, and processed each of these subsets with Binner (v 1.0.0) ^122^. Binner compares metabolite features “binned” by rt and calls putative adduct relationships between these features based on a table of possible adducts, feature *m/z*, and correlation of peak values across samples. Data were split by treatment to avoid possibly conflating signals for different compounds found in only one of the treatments as adducts of each other. Data were natural log transformed, no additional deisotoping step was preformed, the mass tolerance was set to 0.005 Da (biogenic amines and lipids) or 0.003 Da (polyphenols), and default settings were used elsewhere. For biogenic amines and polyphenols, we searched for adducts used previously used by Binner ^122^. For lipids, we supplemented this adduct table with an additional adduct set publicly available online ^123^. We then reunited data subsets to create feature lists that included all peaks that were labeled as primary ions or remained unannotated in either treatment group. In effect, this step only removed peaks that were determined to be putative adduct peaks in both treatments, resulting in the final processed datasets for analysis of: 2955 biogenic amines (90% of QC filtered peaks remained), 4315 polyphenols (88% of QC filtered peaks), and 4924 lipids (78% of QC filtered peaks) (Supplementary Data S1).

### Metabolomic data analysis & feature selection

To select a single set of top-features of interest combined across all three datasets, we applied three statistical approaches (Fig. 2A). For features with values equal to zero, we imputed zeros as 20% of the minimum value measured for that feature. Each analysis we preformed was done independently per LC-MS/MS dataset (biogenic amines, lipids, and polyphenols). We determined if a feature represented a differentially abundant metabolite (DAM) with a t-test (corrected FDR, p ≤ 0.05) and minimum ±1.5-fold change in abundance between sham and infection treatments. Principal component analyses always indicated that only PC1 separated samples by treatment type (see Results). As such, we noted features ranked in the upper 50% of PC1 absolute loading values. Given that our choice of threshold was defined as half of the data, we did not rely on this test solely as merit for discussing a feature, but rather as a way to add confidence to results of other analyses. Both t-tests and PCAs were performed through the MetaboAnalyst interface (v 5.0) ^28^. Finally, we selected features “confirmed” as “important” by Boruta (v 7.0.0, R package) random-forest based analyses with 900 maxRuns and 70,000 ntree settings ^27^. Boruta selects “important” features by z-scores of mean decrease accuracy in decision trees in comparison to randomized versions of the data ^27^. This offers an improvement over standard random-forest analyses for high dimensional data such as in metabolomics, increasing the stability and robustness of the features selected ^124^.

Additionally, we created co-abundance network modules of metabolite features per LC-MS/MS dataset using the WGCNA method ^46^ (Fig. 2A). Although WGCNA was developed primarily for gene expression data, this technique has been used and evaluated for metabolomic data as well ^42, 125, 126^. Heightened activity in a metabolic pathway could lead to many compounds increasing. But simultaneously, their precursors may become depleted. Therefore, correlations within a network could be both positive and negative. As such, we constructed unsigned WGCNA modules. All WGCNA modules were constructed on feature data without imputing zeros, based on the biweight midcorrelation to compare between treatment types (e.g., corFnc=“bicor”, robustY=F, maxPOutlier=0.05) (R package WGCNA, v 1.67). Soft power thresholds were selected according to program recommendations as follows: biogenic amines – 3, polyphenols – 10, and lipids – 11.

We analyzed different groups of selected features for metabolic pathway enrichments. We formed feature sets that combined data across all three LC-MS/MS datasets based on individual top-feature selection analyses (DAM t-tests, PCAs, and Boruta) or their presence only in manipulation samples. Dataset-specific network modules were only analyzed with background features from the LC-MS/MS dataset they belonged to (i.e., amines, polyphenols, or lipids) and if the module was correlated to treatment (Student correlation ≥ |0.5|, p ≤ 0.05) (Fig. 2A).

We preformed Mummichog analyses on all the above feature sets ^30^. This process matched 1,523 unique metabolite features to 6,725 KEGG annotated compounds. For enrichment analyses combining features across datasets, we additionally used a technique based on GSEA ranking features by t-scores returned by t-tests ^127^. In network module pathway enrichment analyses, only Mummichog was used as it sets a hard threshold to define a group of interest versus the background. Membership within or out of a network module was simply defined and could be used for this division. All enrichment analyses were conducted through the MetaboAnalyst interface with the following settings: mixed ion mode, 3 ppm mass tolerance, retention time included, primary ions not enforced, all adducts tested, default currency metabolites, and conducted once for both the *Drosophila melanogaster* and *Saccharomyces cerevisiae* KEGG metabolic pathway and compound databases (Fig. 2A) ^28^. As we sought to characterize the composite metabolome of manipulated ant heads containing both insect and fungal tissue, we chose both a well-studied insect and fungal model organism to improve our odds of matching annotations. We did not use results from one database or the other to discriminate between a host (insect) or parasite (fungus) signal. Neither of the databases would represent the full metabolic repertoire of their respective model organisms and combining them could offer more complete data for comparison. Furthermore, we wished to consider the possibility that due to the relationship of the host and parasite, either organism may be producing metabolites canonically associated with the other as part of its strategy in this antagonism.

We selected significantly enriched pathways based on one of the hypergeometric enrichment results reported by MetaboAnalyst. When using Mummichog only (WGCNA module feature sets), we used the gamma-null adjusted p-value (≤ 0.05). When including a GSEA as a joint analysis (top-features or features present in manipulation only), we used the combined p-value (≤ 0.05), calculated with MetaboAnalyst using the Fisher method. Of the overrepresented pathways selected by the enrichment analyses (p ≤ 0.05), we further imposed thresholds for: (*i*) a minimum of three selected features in the pathway, (*ii*) 1.25-fold enrichment, and (*iii*) three differentially expressed genes (DEGs) (see below) encoding associated enzymes to discard weak signals that would likely be difficult to interpret biologically in a multiomic pathway-level context (Fig. 2A). For cases when a pathway was selected using both *D. melanogaster* and *S. cerevisiae* KEGG databases, a mean p-value and fold-enrichment is reported.

### Multiomics integration of genomic and transcriptomic data

We interpreted our LC-MS/MS data in light of previously collected transcriptomic data when possible ^24^. This RNAseq data was collected with a similar experimental framework, including comparisons of gene expression of both *C. floridanus* and *O. camponoti-floridani* between a control (uninfected ant or fungal culture) and manipulated ant samples. Identification of DEGs (Cuffdiff analysis corrected FDR p ≤ 0.05, minimum two-fold change, minimum 4 RPKM) and relevant statistics are reported in Will *et al.* 2020 ^128^. For each enriched metabolic pathway, we retrieved gene sequence data for enzymes associated with that pathway in KEGG (by enzyme nomenclature EC number) using KEGGREST (v 1.30.1) ^129^. For each enzyme EC, we extracted the first representative gene listed for *D. melanogaster* and *S. cerevisiae*. We searched for reciprocal best matches to those genes in both the *C. floridanus* and *O. camponoti-floridani* genomes using Proteinortho (v 5) with default settings ^130^. While this non-exhaustive approach would not identify every possible *C. floridanus* or *O. camponoti-floridani* gene with homology to known enzymes of interest, it did assist with initially selecting pathway enrichments for deeper analysis. Per pathway, we required that at least three DEGs combined from both transcriptomes of the ant and fungus had homologs to the enzymes associated with that pathway. This selection criterion followed the pathway-level thinking to require at least three contributing metabolite features matched by Mummichog to consider an enrichment result for discussion. Metabolic pathway relationships were based on KEGG data and supplemented by other sources where cited.

## Data Availability

Genomic and transcriptomic data referenced here are available in Will *et al.* 2020 and the GenBank accessions reported therein. Processed metabolomic data are given with this publication as supplementary information. Raw LC-MS/MS peak data have been uploaded to MassIVE (UPLOAD URL PENDING).

## Acknowledgments

We would like to thank Sophia Vermeulen for assisting with testing fungal strain OJ2.1 and Biplabendu Das for their help with ant infections and observations. We also thank Devin Burris for help with ant monitoring. Jordan Dowell provided insightful discussions on LC-MS/MS methods and interpretation for which we are grateful. The Genetics Society of America provided travel support for dissemination of early results of this work. Research funding for this project came from NSF (NSF-CAREER IOS-1941546 awarded to CdB).

## Author Contributions

CdB and IW conceived the study with methodological guidance from GMA. IW performed the experiment and analyses. All authors contributed to interpretation, writing, and revision and have approved this manuscript for submission.

## Additional Information

### Competing Interests

The authors declare no competing interests.

### Main text figure and table legends

**Supplementary Data 1.** LC-MS/MS feature data used for analysis.

**Supplementary Data 2.** KEGG pathway enrichments and annotated features per feature group.

**Supplementary Data 3.** Internal standards used by the WCMC.

**Supplementary Data 4.** Original WCMC data files.

**Supplementary Figure 1.** Acetylcholine and acetylcholinesterase data.

**Supplementary Table 1.** Number of significant features per statistical analysis.

**Supplementary Discussion 1.** Metabolic pathway DEGs.

**Supplementary Discussion 2.** Drug metabolism pathway enrichments.

**Supplementary Discussion 3.** Beta-alanine pathway enrichment.

**Supplementary Discussion 4.** Vitamin B6 metabolism.

**Supplementary Discussion 5.** Serotonin and octopamine metabolism.

**Supplementary Discussion 6.** Glutamine and GABA metabolism.

**Supplementary Methods.** Expanded LC-MS/MS methods.

## Notes

### Competing Interest Statement

The authors have declared no competing interest.

### Summary of Updates

Streamlining of the writing and reporting. Figure updates for clarity.

