## Supplemental Information for "Multiomic interpretation of fungus-infected ant metabolomes during manipulated summit disease"

**Supplementary Table 1****. Number of significant features per analysis type.**

We performed three analyses (DAMs, Boruta, and PCA) to detect features that significantly changed in their abundance in manipulated ant heads as compared to healthy ant heads. Among the DAMs that we detected, most increased in their abundance during manipulation in the biogenic amines and lipids datasets, whereas the polyphenol DAMs slightly more often decreased. In addition to determining DAMs and Boruta confirmation, we also selected features in top 50% of PC1 from PCAs. These statistics are not reported in the table as they are half of the total features per dataset by definition. Top features that passed at least two of these selection analyses were further used in enrichment analyses, placed in the context of infection and behavioral manipulation.

| **Dataset** | **Total features** | **Total DAMs** | **Increased DAMs** | **Decreased DAMs** | **Boruta confirmed** | **Top**  **features** |
| --- | --- | --- | --- | --- | --- | --- |
| Biogenic amines | 2955 | 1774 | 1129 | 645 | 1226 | 1535 |
| Polyphenols | 4315 | 2641 | 1141 | 1500 | 1619 | 2262 |
| Lipids | 4924 | 2995 | 2390 | 605 | 2477 | 2671 |
| Total | 12194 | 7410 | 4660 | 2750 | 5322 | 6468 |

**Supplementary Discussion S1. Metabolic pathway DEGs were mostly downregulated.**

Among the enzyme homologs used to select pathway enrichments across all feature sets, most ant and fungal DEGs were downregulated from the healthy ants to manipulation (ant: 142 of 157, 90%, fungus: 166 of 268, 62%). In both cases, these numbers of downregulated DEGs were much greater than the transcriptome-wide DEG ratios during manipulation for ant (72% downregulated) and fungus (41% downregulated) (binomial exact test, p = 2E-8 and 6E-12, respectively). The metabolic pathway DEG estimates are based on a coarse analysis using homology to a constrained subset of well-annotated fly and yeast enzymes listed in KEGG; however, this sample of genes could plausibly be a proxy for broader patterns among metabolism-related DEGs. In *Camponotus* hosts, the stark downregulation of genes related to these metabolic pathways may indicate a general slowing of metabolism in a weakened host, terminally infected and manipulated in the last hours of life. In the fungal parasite, the significant reduction of gene expression of these pathways could more plausibly reflect a state of *O. camponoti-floridani* metabolism tuned to late infection and manipulation processes. By the time of manipulation, the fungus has been engaged with host immunity and modulation of behavior and has access to host metabolites to scavenge. Genes related to those functions may be upregulated while genes related to non-critical metabolic pathways may be downregulated. Furthermore, many upregulated fungal genes important for manipulation may not produce enzymes involved in major metabolic pathways, but rather extracellular effectors (e.g., involved in protein-protein interactions), post-translational regulators, and others.

**Supplementary Discussion S2. Drug metabolism pathway enrichments hint at possible detoxification activity during infection and manipulation.**

Enriched pathway “drug metabolism – other enzymes” in the top features and module A1 (p = 0.030 and 0.006, fold-enrichment = 1.5 and 2.1, respectively) was complemented by the other pathway enrichments “drug metabolism – P450” in network modules L3 and P3 (p = 6.7E-4 and 0.024, fold-enrichment = 10.0 and 1.4, respectively) and “metabolism of xenobiotics by P450” in A1 and P2 (p = 0.012 and 0.29, fold-enrichment = 1.6 and 1.5, respectively). Each of these KEGG pathway annotations contain multiple distinct metabolic pathways pertaining to the detoxification metabolism specific drugs or compounds. Altered drug metabolism pathways could reflect antagonist host-pathogen interactions employing metabolites with similar structure as the drugs and toxins specifically described in the KEGG pathways. Our data had no metabolite features annotated with compounds found in these pathways. Therefore, we retrieved Mummichog matches between “empirical compounds” and KEGG compounds with R package keggrest (v 1.34.0) for discussion ^1^. Mummichog is not a feature annotation tool meant to confidently pair a feature to a single best annotation, but we used these annotations to look for possible patterns within the metabolic pathways in lieu of more concrete data. Across all three pathways, matched compounds were diffusely populated across multiple drug-specific sub-pathways and in all cases either reduced in abundance or not significantly changed. Given this trend we cannot offer compound-specific biological interpretations for these pathway enrichments or why such metabolites would be found reduced during manipulation.

However, outside of these network modules enriched for drug and xenobiotic metabolism, an increased DAM matched to aflatoxin B1-exo-8,9-epoxide by Mummichog as its only compound match. This feature was only detected in manipulated samples (p = 2.4E-14, median peak value = 749,741, top feature). We single out this compound match as it is a derivative of aflatoxin B1, a well-recognized mycotoxin produced by fungi with insecticidal properties ^2,3^. Two other features matched aflatoxin B1-exo-8,9-epoxide, one a decreased DAM and the other not differentially abundant. Given that the increased DAM was only detected in manipulated ants, it appears as the most sensible feature match to a degradation product of a specialized fungal metabolite. In the *O. camponoti-floridani* genome, multiple genes putatively related to mycotoxin and secondary metabolite production were identified and many were upregulated during manipulation ^4^. Whether this putative aflatoxin derivative reflects hypothesized fungal toxin production during infection of the ant host remains unclear.

**Supplementary Discussion S3. Beta-alanine metabolism may lead to carnosine production that could alleviate the physiological stress of manipulation and infection on host ants.**

Within network module P2 and top features, we detected an enrichment signal for the “beta-alanine metabolism” pathway (p = 0.025 and 0.002, fold-enrichment = 1.8 and 1.5, respectively). Two amino acids participating in this pathway were reduced DAMs in manipulated ants, β-alanine (p = 0.012, 1.6-fold decrease) and histidine (p = 1.3E-5, 4.5-fold decrease, top feature). We also observed increased carnosine (p = 6.4E-10, 4.1-fold increase, top feature), which is a dipeptide formed from β-alanine and histidine. Carnosine and its analogs are primarily known from vertebrates, with rare observations in mollusks ^5,6^. Indeed, within healthy ant samples, the signal for carnosine was detected at nearly the same levels of our mock blank-collection samples (only 1.2-fold higher). In vertebrates, carnosine is found largely in muscle and the olfactory bulb where it has protective effects for both muscular and neuronal function by possible pH buffering and antioxidant properties ^7–10^. Neither the *O. camponoti-floridani* nor *C. floridanus* genomes had robust homology to carnosine synthase genes that encode the enzyme for the formation of carnosine. However, the fungal genome contains more likely candidates. For protein homology searches we used BLASTp with default settings ^11^. For protein sequence alignment, we used the MAFFT plugin (v 1.4.0, Biomatters) through Geneious (v 2022.0.2, Biomatters) with default settings ^12^.

Carnosine synthases possibly originated from an ancestral gene containing two ATP-grasp domains and a C-terminal D-alanine-D-alanine ligase domain ^13^. In the fungal parasite, there are seven genes containing both of these domains, four of which include the two ATP-grasp and one D-alanine-alanine ligase composition. No ant genes include both domains. However, a lack of strong homology between these fungal genes and animal carnosine synthases leaves the mechanism of production of carnosine in *C. floridanus* manipulated by *O. camponoti-floridani* an open question. We also cannot discount a possible endosymbiont or microbiome resident being involved in this process. Regardless of which organism produced carnosine, it may use β-alanine and histidine to form carnosine as a protective agent against the physiological stresses of hyperactive muscular activity (increased locomotion, tremors, biting) and/or buffer the nervous system against excessive damage until the host has been successfully behaviorally manipulated.

**Supplementary Discussion S4. Vitamin B6 metabolism may change during infection and impact activity the AADC enzyme needed for dopamine synthesis.**

One of the many functions of vitamin B6 is as a co-factor for AADC and therefore it may participate in the modulation of dopamine and serotonin production. Largely, insects cannot synthesize active vitamin B6 *de novo* but rather metabolize B6 vitamers obtained by diet and symbiotic microbes to meet their vitamin B6 demands ^14,15^. Network module A5 had two compound annotations that specifically suggest changes in vitamin B6 metabolism. Pyridoxamine is a vitamer form of vitamin B6 and was annotated in network module A5 as an increased DAM (p = 4E-7, 3.5-fold increase, top feature). Pyridoxamine is not the active B6 vitamer used as a co-factor for AADC, but is converted to the active pyridoxal 5’-phosphate vitamer by two additional enzymatic steps ^16^. Negatively correlated to pyridoxamine in network module A5, glutamine, one of three substrate molecules needed for the fungus to form active vitamin B6 *de novo* ^17^, was reduced during manipulation (p = 7E-6, 3.3-fold decrease, top feature). Similarly, L-glutamine was also detected in network module P2 as a DAM reduced in abundance (p = 8E-10, 3.9-fold decrease, top feature). Both modules A5 and P2 contained metabolomic signals relevant to dopamine metabolism.

In *O. camponoti-floridani*, putative homologs to yeast SNZ-1 (synthase domain) and SNO-1 (glutaminase domain) genes, which encode proteins that come together to catalyze *de novo* synthesis of active vitamin B6 ^18^, showed mixed DEG profiles during manipulation. The SNZ-1-like gene was downregulated while the SNO-1-like gene did not significantly change in expression. If vitamin B6 synthesis was reduced in the fungus, one might predict that glutamine levels would increase, which we did not find. This reduction of glutamine could be related to other biological processes, such as glutamate metabolism (Supplementary Discussion 6). Consistent with insects lacking a *de novo* pathway ^15^, no homologs to these genes were found in the ant genome, nor were any *D. melanogaster* representatives of this enzyme group listed in the KEGG database. In *O. camponoti-floridani*, a putative homolog of the yeast PDX3 gene (a pyridoxamine 5'-phosphate oxidase) that converts a pyridoxamine 5’-phosphate intermediate to active vitamin B6 was also downregulated during manipulation. Taken together, the fungus may be scavenging host vitamin B6 and reducing its own synthesis of active vitamin B6 as evidenced by downregulated SNZ-1 and PDX3-like genes and an accumulation of the precursor vitamer pyridoxamine. Possibly, the accumulating pyridoxamine is available to the ant host, and her sustained expression of a pyridoxamine 5'-phosphate oxidase homolog (not a DEG, albeit with a 1.4-fold increase) continues to produce active vitamin B6. Speculatively, if substrate availability limited ant synthesis of active vitamin B6, the availability of pyridoxamine could even lead to elevated vitamin B6 levels. In scenarios where active vitamin B6 is more abundant in ant tissues, it may be working in concert with elevated AADC to substantially increase dopamine production. Alternatively, if vitamin B6 is scarce, upregulated AADC could represent a homeostatic response to insufficient dopamine production, at least in part.

**Supplementary Discussion S5. Serotonin and octopamine biosynthesis may accompany changes in dopamine metabolism to affect host behavior.**

*Altered serotonin biosynthesis during manipulation could be linked to manipulated behaviors.* In consideration of the proposed changes in dopamine metabolism, which is linked to serotonin biosynthesis by the shared upregulated AADC enzyme, we additionally searched our datasets for signals tied to serotonin metabolism. Tryptophan can be converted to serotonin by two enzymatic steps (the second of which is mediated by AADC). In polyphenol network module P2, which contained L-tyrosine, L-DOPA, and the tyrosine and phenylalanine metabolism pathway enrichments discussed as elements of dopamine metabolism, we detected a tryptophan signal that was 2.2-fold reduced (selected by Boruta but not a DAM, p = 0.283). This tryptophan signal was supported by an annotated tryptophan adduct that was similarly 2.1-fold reduced and selected by Boruta but not our DAM analysis. In the amine dataset, tryptophan was also detected in network module A5, which carried the phenylalanine feature annotation. This measurement of tryptophan was not selected by any analysis (1.3-fold increase during manipulation). Given the conflicting signals, we cannot offer a concrete conclusion on the relative abundance of tryptophan during manipulation.

However, in light of the two decreased tryptophan features from the polyphenol dataset, the upregulation of AADC, and the plausible behavioral link to changing serotonin levels during infection, we propose here a hypothetical scenario of increasing serotonin levels. Upregulation of ant AADC could indicate increased serotonin synthesis and the simultaneous depletion of tryptophan would be consistent with ongoing metabolism of this compound providing the precursor to serotonin. Furthermore, a stably abundant feature annotated as 3-hydroxykynuerine (a tryptophan derived compound) could indicate that the reduction of tryptophan does not reflect widespread increasing metabolism in a competing non-serotonergic pathway. We detected two features annotated as 3-hydroxykynuerine that were not DAMs nor correlated with other features in network modules. Although, tryptophan can be metabolized into many possible compounds, upwards of 95% of tryptophan is metabolized in the kynurenine degradation pathway in mammals, which can have immune- and neuromodulatory effects ^19,20^. Further supporting that there was no broad increase of kynurenine pathway metabolism, expression of the rate-limiting enzyme to enter this direction of tryptophan metabolism is constant in the ant and downregulated in the fungus (indoleamine 2,3-dioxygenase in the fungus). Genes for kynurenine 3-monooxygenase that catalyzes the production of 3-hydroxykynuerine were also not DEGs in either organism. However, we did find two upregulated genes encoding enzymes involved in kynurenine and kynurenic acid synthesis. The host upregulated a putative kynurenine/alpha-aminoadipate aminotransferase that catalyzes synthesis of kynurenic acid and xanthurenic acid. Kynurenic acid particularly can have neuroprotectant effects, but has also been associated to neurological disease ^20–24^. Additionally,

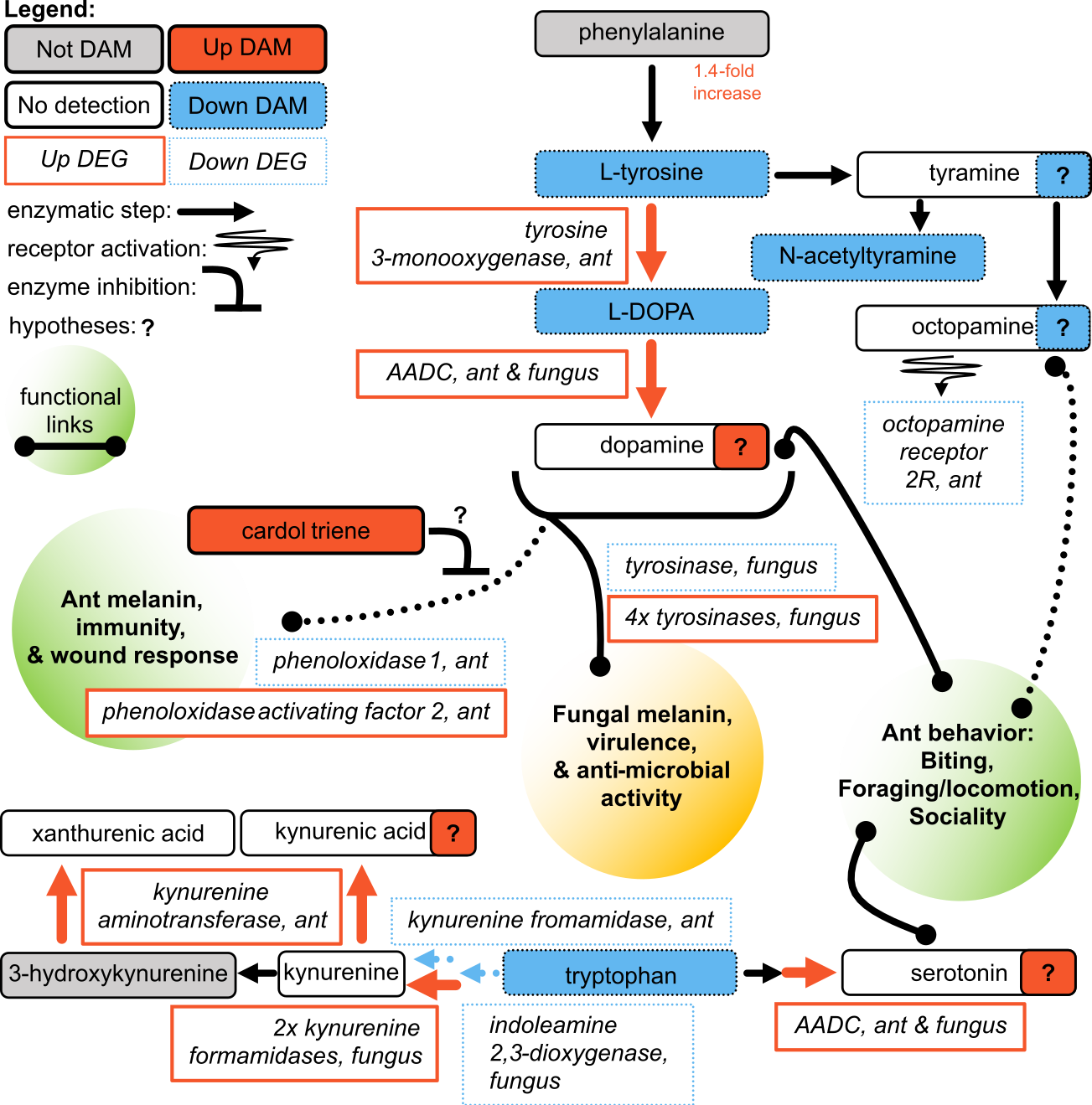
**Increasing serotonin and decreasing octopamine may accompany the hypothesized increase in dopamine.** Kynurenic acid may also be increasing and acting as a neuroprotectant during manipulation. L-amino acid decarboxylase (AADC) catalyzes the final step of dopamine and serotonin synthesis and is upregulated by manipulated ants. A reduction of L-DOPA and tryptophan suggest that these compounds are being converted into dopamine and serotonin, respectively. Upregulation of an ant kynurenine aminotransferase possibly increases or maintains (if the kynurenine precursor is reduced) kynurenic acid levels. The possible of reduction of octopamine concentration follows the reasoning that it requires L-tyrosine for its synthesis, which is reduced during manipulation, and there is no compensatory upregulation of enzyme genes in the octopamine pathway to compete with upregulated dopamine biosynthesis. Increases are indicated by solid lines and red color, decreases by dotted lines and blue color, metabolites are in rounded rectangles, genes are in rectangles, annotated metabolites that were not DAMs have gray color, compounds with white color were not identified in the LC-MS/MS data, and hypothesized functional links are given in the shaded circles. Metabolic steps that did not have DEG are shown as black arrows when useful for context but are omitted elsewhere. A squiggled arrow indicates metabolite interaction with a receptor. Tryptophan is given colored as a reduced abundance DAM, but was selected only by the Boruta analysis only and not the differential abundance t-test. Additionally, a second metabolite feature was putatively identified as tryptophan and not selected by any analysis, but lacked a supporting adduct feature, which the Boruta-selected feature did have.

*Ophiocordyceps* upregulated two kynurenine formamidases, the much more highly expressed of which we predicted to be secreted. The fungus may be using this enzyme exported to the host environment to promote synthesis of kynurenine, the precursor for kynurenine aminotransferase mediated synthesis of kynurenic acid. In contrast, *Camponotus* downregulated a kynurenine formamidase homolog. Taken together, we infer that reduced tryptophan abundance represents a metabolic shift towards serotonin synthesis rather than broadly upregulating the alternative pathway of kynurenine metabolism; but certain key elements of the kynurenine pathway may become more active during manipulation without drawing more heavily on available tryptophan.

*Mixed signals in octopamine metabolism and reception indicated a possible reduction of octopamine during manipulation.* With data supporting changing dopamine and serotonin metabolism, we also investigated octopamine metabolism, which shares a common precursor with dopamine through L-tyrosine. In addition to the proposed L-tyrosine depletion feeding into dopamine synthesis, L-tyrosine can also be converted to tyramine – the precursor to octopamine. However, unchanging gene expression in the ant for tyrosine decarboxylase indicates this was not happening at an accelerated pace. Possibly, upregulation of the rate-limiting tyrosine 3-monooxygenase, which converts L-tyrosine to L-DOPA, outcompeted the tyramine and octopamine pathway for L-tyrosine substrate. In line with this scenario, we did observe reduction of N-acetyltyramine, a product synthesized from tyramine. Arylalkylamine N-acetyltransferases, and specifically in *D. melanogaster*, dopamine N-acetyltransferase converts tyramine, dopamine, and serotonin to N-acetylated forms, generally along melanin pathways related to immunity, cuticle sclerotization, or melatonin ^25–28^. The putative ant homolog to dopamine N-acetyltransferase was not a DEG, further suggesting that tyramine nor these other neurotransmitters were converted to non-neurotransmitter acetylated forms more than in healthy individuals. Given that levels of N-acetyltyramine were reduced but the enzyme for this conversion did not change in expression suggested that either tyramine was being depleted for reasons other than N-acetyltyramine synthesis, or, N-acetlyyramine was being consumed at an accelerated pace. As the gene for conversion of tyramine to octopamine was not differentially expressed in the ant, it appears plausible that levels of octopamine would have also decreased if the availability of tyramine was reduced. However, the ant significantly downregulated a gene putatively encoding octopamine receptor 2, which could be more reflective of a homeostatic response to accumulating octopamine levels. In consideration of the reduced L-tyrosine and N-acetyltyramine levels, with upregulated tyrosine 3-monooxygenase, we propose a reduction in octopamine as the more likely scenario. If so, the possibility exists that the downregulation of the octopamine receptor could be due to direct fungal interference with gene regulation.

**Supplementary Discussion S6. Glutamine and GABA in the context of ant neuromodulation and fungal growth metabolism.**

In insects and other animals, glutamate and GABA are well documented neuromodulators with notable roles at the insect neuromuscular junction, among their other metabolic functions ^29–34^. In *C. floridanus*, multiple genes putatively encoding receptors containing domains associated with GABA or glutamate response were negatively correlated to manipulation by a two WGCNA gene modules ^4^. Additionally, a putative biosynthetic gene cluster for an aflatrem-like compound in *O. camponoti-floridani* displayed a strong upregulation during manipulation of host ants ^4^. Aflatrem has been shown to cause tremors, confusion, and changes in activity levels in mammals by mechanisms including disruption of GABAergic and glutamatergic neuronal activity ^35–38^. However, these possible disruptions of ant neuronal function, do not appear to be clearly tied with dramatic changes in compound abundances or metabolic enzymes. Glutamic acid (the conjugate acid of glutamate) did not qualify as a DAM, although it displayed marginal reduction in concentration short of our 1.5-fold change threshold (p = 0.007, 1.4-fold decrease) (Fig. 19). This glutamate feature was grouped in network module A5. Also in A5, we detected changes in multiple DAMs with biosynthetic links to glutamate and GABA. Directly interconvertible with glutamate, we detected reduced glutamine (p = 1.8E-6 and 6.2E-11, 3.3- and 3.8-fold decrease, in network modules A5 and P2, respectively) and N-acetylglutamic acid (p = 0.022 and 7.3E-6, 2.0- and 1.8-fold decrease, in network modules A5 and P18, respectively). Within two or three biosynthetic steps from glutamate or GABA, we found increased succinate (p =0.002, 2.1-fold increase).. At a similar biosynthetic distance, we detected reduced ornithine (p = 3.1E-5, 2.2-fold decrease), proline (p = 0.001, 1.9-fold decrease), putrescine (p = 0.001, 1.9-fold decrease), and arginine (p = 1.4E-9 and 2.1E-7, 2.7- and 2.3-fold decrease, both confirmed by Boruta), with an additional arginine annotation that was not selected (p = 0.005, 1.3-fold decrease only).

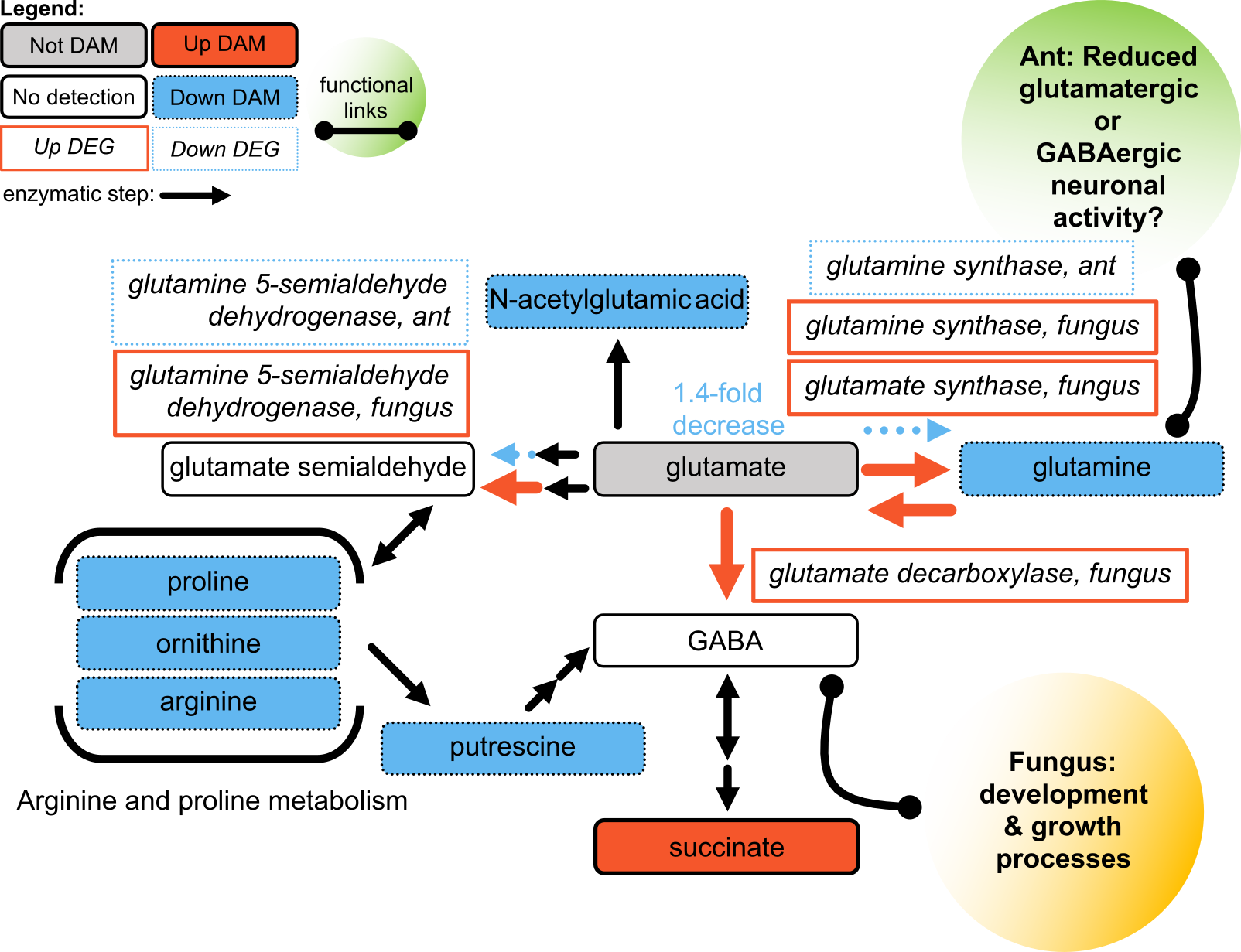

**Changes in glutamate and GABA metabolism may change with different effects for host and parasite.** Possible increased GABA biosynthesis in the fungus that may be used for growth and development metabolism; while a wide-spread reduction of these compounds in the manipulated ant could relate to neuronal and behavioral activity. Legend as Supplementary Discussion S5.

Cytidine has been associated with reduced glutamate and glutamine with anti-depressive neurological effects in certain contexts ^39^. Cytidine was an increased DAM during manipulation that was only detected reliably in manipulated ant samples (p = 7.4E-8, 19.8-fold increase, top feature). To what degree this elevated cytidine content is available to host cells or is an aspect of the proliferating fungal cell population (e.g., nucleic acid metabolism) is unclear. Cytidine can also be interconverted with uridine, an increased DAM in manipulated ants (p = 3.0E-7, 5.4-fold increase, top feature), which in combination with choline has shown some neuroprotective effects, likely related to phosphatidylcholine synthesis ^40^.

Furthermore, in *Camponotus*, a putative glutamine synthase (converting glutamate to glutamine) was downregulated during manipulation. Ant glutamate synthase (reversely converting glutamine to glutamate) and a glutamate decarboxylase (converting glutamate to GABA) were not DEGs but had marginal 1.5-fold reductions. Glutamate decarboxylase is also an enzyme requiring vitamin B6 as a co-factor ^41,42^, similar to AADC (see above). Ant enzymes associated with the GABA shunt pathway to convert GABA to citrate cycle derivatives (e.g. succinate) were also not DEGs ^43^. Glutamate semialdehyde can also be interconverted between glutamate and compounds that have become scarcer during manipulation – arginine, ornithine, and proline. Network modules A1 and P2 were enriched for the “arginine and proline metabolism” pathway related to these compounds (p = 0.013 and 0.039, 1.5- and 1.9-fold reduced, respectively). Taken together, in the ant, there appears to be a general holding or slight reduction in biosynthetic activity and decreased compound abundances related to glutamate and GABA. Although, this could have physiological, neuronal, and behavioral repercussions in the ant – the metabolomic signals could also indicate fungal metabolism.

In the fungal transcriptome, multiple putative genes for glutamate-related enzymes were DEGs during manipulation. Among these genes, a putative glutamate decarboxylase, which converts glutamate to GABA, was upregulated. Additionally, both a glutamine synthase and a glutamate synthase were upregulated, suggesting heightened metabolic activity interconverting glutamate and glutamine. Given the reduced levels of glutamine, the mostly consistent levels of glutamate, upregulation of the putative fungal glutamate decarboxylase, and increase of succinate – we suspect the fungus increased GABA production by converting glutamine to glutamate to GABA, which in turn was converted to succinate. Broadly, GABA can be used by various fungi in nitrogen metabolism, conidiation, and development processes ^44^. For example, in *Trichoderma atroviride*, GABA and glutamate decarboxylase appear to be important players in normal fungal growth. Deletion of glutamate decarboxylase and reduction of GABA decreased the growth and germination rates and produced atypical highly-branched hyphal morphologies ^45^. As *O. camponoti-floridani* proliferates in the terminally manipulated host, soon needing to rapidly colonize the host cadaver in a hyphal growth stage, GABA-dependent growth processes may be already upregulated at the time of manipulation, a few hours before host death. The fungus may also be using an upregulated putative glutamine 5-semialdehyde dehydrogenase that catalyzes one of two steps to convert glutamate into glutamate semialdehyde, which in turn can be used to synthesize amino acids that were reduced in manipulated ants. Meanwhile, the ant downregulated a homolog to this gene.

Glutamate and GABA are also important neurotransmitters, insect neuromuscular modulators, and components of metabolism more broadly. Glutamate and many of the compounds that are biosynthetically linked to glutamate and GABA are reduced during manipulation. Manipulated ant gene expression suggests that glutamate conversion into GABA remains steady, while two alternative branches of glutamate metabolism are downregulated. These transcriptomic data possibly signifying increased substrate availability for GABA production or homeostatic attempts to maintain glutamate levels amid widespread depletion of glutamine and other compounds. The fungus meanwhile appeared to be increasing consumption of glutamate, including by upregulation of a gene putatively encoding a glutamate decarboxylase to produce GABA. We consider it most likely that the fungus is using GABA and glutamate derivatives for its own metabolic demands during growth and development. Although, we cannot discount it exporting GABA into the host environment.

**Supplementary Figure 1.**

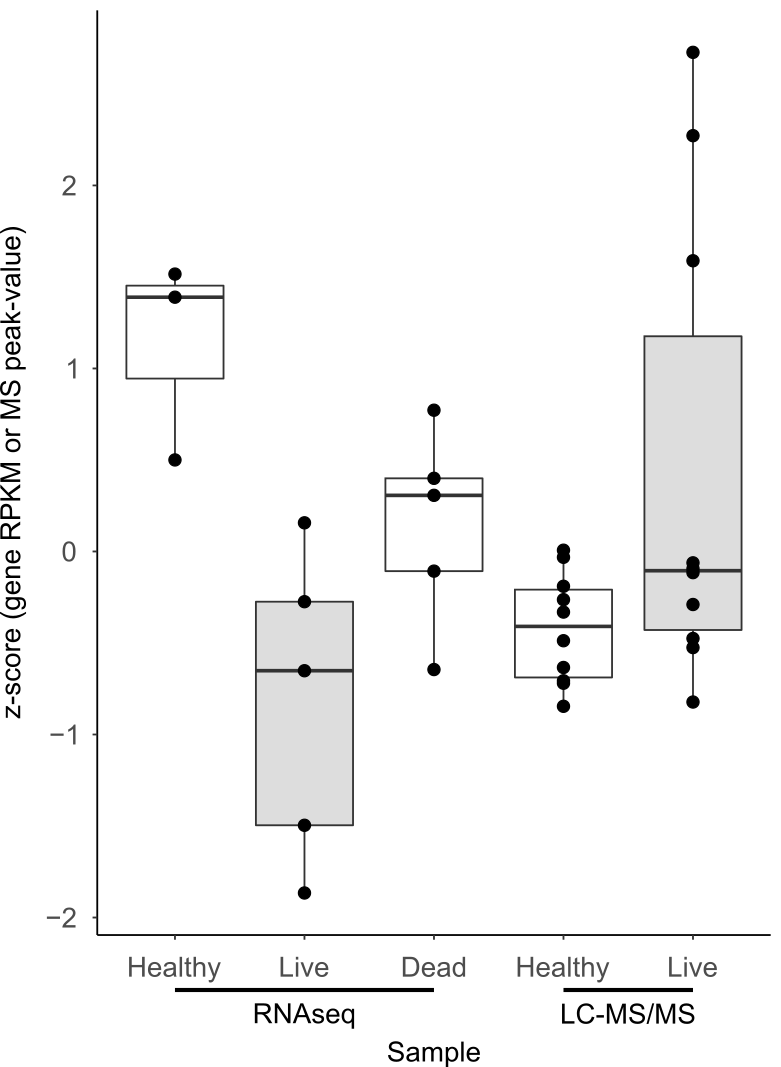

**Supplementary Figure 1. Comparison of acetylcholinestaerase-2 gene expression by *C. floridanus* and acetylcholine LC-MS/MS peak values.** In both the previous transcriptomic and current metabolomic works, live manipulated ants showed the greatest variation in acetylcholine-related signals (gray). Dead samples refer to moribund manipulated ants that were collected as soon as they become unresponsive and presumably dead, or nearly so. Although precise sampling at a fine timescale and increased replication are required to further investigate possible variation within the live manipulation category – the fast action of acetylcholinesterase to reduce acetylcholine would be in line with the proposed “high” and “low” acetylcholine sub-grouping. Data are shown as z-scores scaled according to their data type (gene expression or peak values).

**Supplementary Methods.**

Biogenic amine HILIC LC-Q-TOF in electrospray ionization (ESI) positive and negative mode materials and methods were as follows: Agilent 6530 Q-TOF (ESI positive) and Agilent 6550 Q-TOF (ESI negative) spectrophotometers; Waters Acquity UPLC BEH Amide VanGuard (1.7 μm, 2.1 mm × 5 mm) pre-column; Waters Acquity UPLC BEH Amide (1.7 μm, 2.1 mm × 5 mm) column; 40 C° column temperature; ultrapure water with 10 Mm ammonium formiate, 0.125% formic acid, pH 3 mobile phase (A); acetonitrile:ultrapure water 95:5, with 10 mM ammonium formiate, 0.125% formic acid, pH 3 mobile phase (B); 1 μL injection volume; 4 C° injection temperature; 0.4 mL/min flow rate; 0 min 100% (B), 0–2 min 100% (B), 2–7 min 70% (B), 7.7–9 min 40% (B), 9.5–10.25 min 30% (B), 10.25–12.75 min 100% (B), 16.75 min 100% (B) elution gradient; 4.5 kV capillary voltage; 45 eV collision energy; 3 Da precursor isolation width; 60 – 1200 Da scan range; 2 spectra/sec spectral acquisition speed; 10,000 mass resolution.

Lipid CSH LC-Q-TOF in ESI positive and negative mode materials and methods were as follows: Agilent 6546 Q-TOF spectrophotometer; Waters Acquity UPLC CSH C18 (1.7 μm, 2.1 mm x 100 mm) column; 65 C° column temperature; acetonitrile:water 60:40, 10 mM ammonium formiate, 0.1% formic acid mobile phase (A); isopropanol:acetonitrile 90:10, 10 mM ammonium formiate, 0.1% formic acid mobile phase (B); 1.7 μL (ESI positive) and 5 μL (ESI negative) injection volume; 4 C° injection temperature; 0.6 mL/min flow rate; 0 min 15% (B), 0–2 min 30% (B), 2–2.5 min 48% (B), 2.5–11 min 82% (B), 11–11.5 min 99% (B), 11.5–12 min 99% (B), 12–12.1 min 15% (B), 12.1–15 min 15% (B) elution gradient; 3.5 kV capillary voltage; 60 – 1200 Da scan range; 10,000 (ESI positive) and 20,000 (ESI negative) mass resolution.

Polyphenol PFP LC-QE in ESI positive and negative mode materials and methods were as follows: Thermo Scientific Q Exactive spectrophotometer; Kinetex PFP column (1.7 μm, 2.1 mm × 30 mm); ultrapure water, 0.1% formic acid mobile phase (A); acetonitrile, 0.1% formic acid mobile phase (B); 5 μL injection volume; 0.4 mL/min flow rate. All QC pools were reported to fail upon injection and therefore not used in data filtering procedures.
